# Wilson disease mechanisms manipulate ATP7B activity

**DOI:** 10.64898/2026.11.16.766959

**Authors:** Tomáš Heger, Orsolya Mozner, Ida M. Skøtt, Thibaud Dieudonné, Anna L. Duncan, Poul Nissen, Charlott Stock

## Abstract

Copper is an essential cofactor for enzymes involved in diverse processes including mitochondrial respiration and the metabolism of neurotransmitters, collagen and melanin. However, its redox activity requires tight homeostatic control. The P-type ATPases ATP7A and ATP7B help deliver copper to cuproenzymes, and ATP7B mediates the excretion of excess copper into bile. Pathogenic *ATP7B* variants cause Wilson disease (WD), leading to copper accumulation in organs including the liver, kidney, and brain, and to disruption of copper-dependent processes. Current treatments primarily reduce systemic copper levels, while emerging gene therapies aim to restore ATP7B function. A complementary therapeutic strategy would be to rescue dysfunctional ATP7B using pharmacological chaperones, but their development is limited by an insufficient understanding of how WD variants perturb ATP7B structure and function. Here, we present a cryo-EM structure of human ATP7B reconstituted in lipid nanodiscs together with coarse-grained molecular dynamics simulations, as well as mutagenesis, functional and sequence analyses. We identify a salt bridge (E754-R969), crucial for the transition between inward- and outward-facing conformations, which is abolished by the R969Q disease variant. Additionally, we find a conserved stabilizing cholesterol-binding site severely disrupted by the WD variant W939C. Furthermore, we present new insights into the dynamics of metal binding domains 5 and 6 hinting at a copper relay mechanism in ATP7B. Together, these findings reveal lipid-and domain-mediated mechanisms of ATP7B regulation and explain how specific WD variants disrupt them, providing a structural framework for mutation-directed therapeutic strategies, including pharmacological chaperones.

## Introduction

Cellular metabolism relies on numerous redox reactions, many of which are catalysed by metalloenzymes containing redox-active metals such as copper or iron. Copper serves as an essential cofactor in enzymes including cytochrome *c* oxidase (complex IV) of the mitochondrial respiratory chain, lysyl oxidase important for connective tissue integrity, monoamine oxidase metabolizing neurotransmitters, tyrosinase producing melanin, or antioxidant enzyme superoxide dismutase^1^. Despite its essential physiological roles, copper becomes toxic when not tightly regulated^2^.

In humans, several transport proteins, including solute carriers and the P-type ATPases ATP7A and ATP7B, maintain copper homeostasis. ATP7A and ATP7B differ in tissue distribution, systemic role in copper handling, and the enzymes they supply with copper. ATP7A mediates copper uptake, and pathogenic variants cause Menkes disease, characterized by copper deficiency in the liver and brain but accumulation in the kidney and intestine^3^. By contrast, ATP7B primarily mediates biliary copper excretion, and pathogenic variants cause Wilson disease (WD)^4^.

WD symptoms arise from copper overload mainly in the liver, brain and kidney, with progressive liver failure, possibly requiring a liver transplant, as one of the most severe manifestations. Other symptoms may include neuropsychiatric diseases, renal tubular dysfunction, or musculoskeletal disorders^4–6^. WD is an autosomal monogenic recessive disorder caused by composite heterozygous genotypes or homozygosity for more common mutations. Consistent genotype-phenotype correlation is made impossible by the manifold permutations of disease-causing mutations, and functional characterization beyond common mutations remains scarce. There are more than 600 known pathogenic variants in the *ATP7B* gene in the ClinVar database^7^ and over 900 variants associated with WD in the Human Gene Mutation Database^8^ (HGMD®; both databases accessed 28 October 2026). WD mutations cover the entire sequence of ATP7B (Suppl. Fig. 1A), suggesting diverse loss-of-function traits. Indeed, the two most common WD mutations R778L (predominant among individuals of East-Asian ancestry, dbSNP^9^ ID: *rs28942074*) and H1069Q (predominant among individuals of European ancestry, dbSNP^9^ ID: *rs76151636*) locate to the cytoplasmic ion entry pathway and the nucleotide-binding domain, respectively, and disrupt ATP7B function via different mechanisms (Suppl. Fig. 1B–C). With the estimated prevalence between 1:30,000 and 1:50,000, and heterozygous carrier frequency between 1:25 and 1:90, the disease burden is likely underestimated^10,11^. However, when recognized early, copper toxicity can be treated via a restrictive diet and small-molecule copper-chelating drugs^12^.

Gene therapies, including gene augmentation and gene editing approaches, could potentially be used to treat WD and are currently undergoing clinical development (ClinicalTrials.gov IDs: NCT07173933, NCT04884815, NCT06663878, NCT06650319, NCT07748403, and terminated NCT04537377). Despite some conflicting results^13^, proof-of-concept studies have demonstrated that manipulating mutant ATP7B can lead to phenotype mildening^14–16^, similar to a cystic fibrosis phenotype when rescuing CFTR function^17,18^. Mechanistic understanding of ATP7B structure-function relationships remains crucial for the interpretation of variants and their potential targeting in therapies.

P-type ATPases like ATP7B generally operate through a so-called Post-Albers cycle with transitions between inward-open E1 and outward-open E2 conformational states interspersed by occluded states^19,20^. ATP binding at the cytoplasmic domains and transport substrate binding in the membrane domain (Cu(I) at a CPC motif for ATP7B) lead to the formation of an occluded E1P phospho-enzyme intermediate. Upon a conformational switch to an outward-oriented E2P state, copper is released and the enzyme undergoes dephosphorylation, facilitated by the internal phosphatase activity of the A-domain (TGEA loop for ATP7B).

Previous studies determined crystal structures of bacterial homologues^21–23^ and cryo-EM structures of frog and inactive human ATP7B^24,25^ and of the rice ortholog HMA4^26^. As a P_IB_-ATPase, ATP7B features two additional transmembrane helices, labelled MA and MB, within its transmembrane M-domain, which otherwise comprises six standard transmembrane segments (M1–M6). The cytoplasmic side of the pump consists of three core domains common to P-type ATPases: the nucleotide-binding (N-) domain, the phosphorylation (P-) domain and the actuator (A-) domain (Figure 1A)^19^.

**Figure 1:**
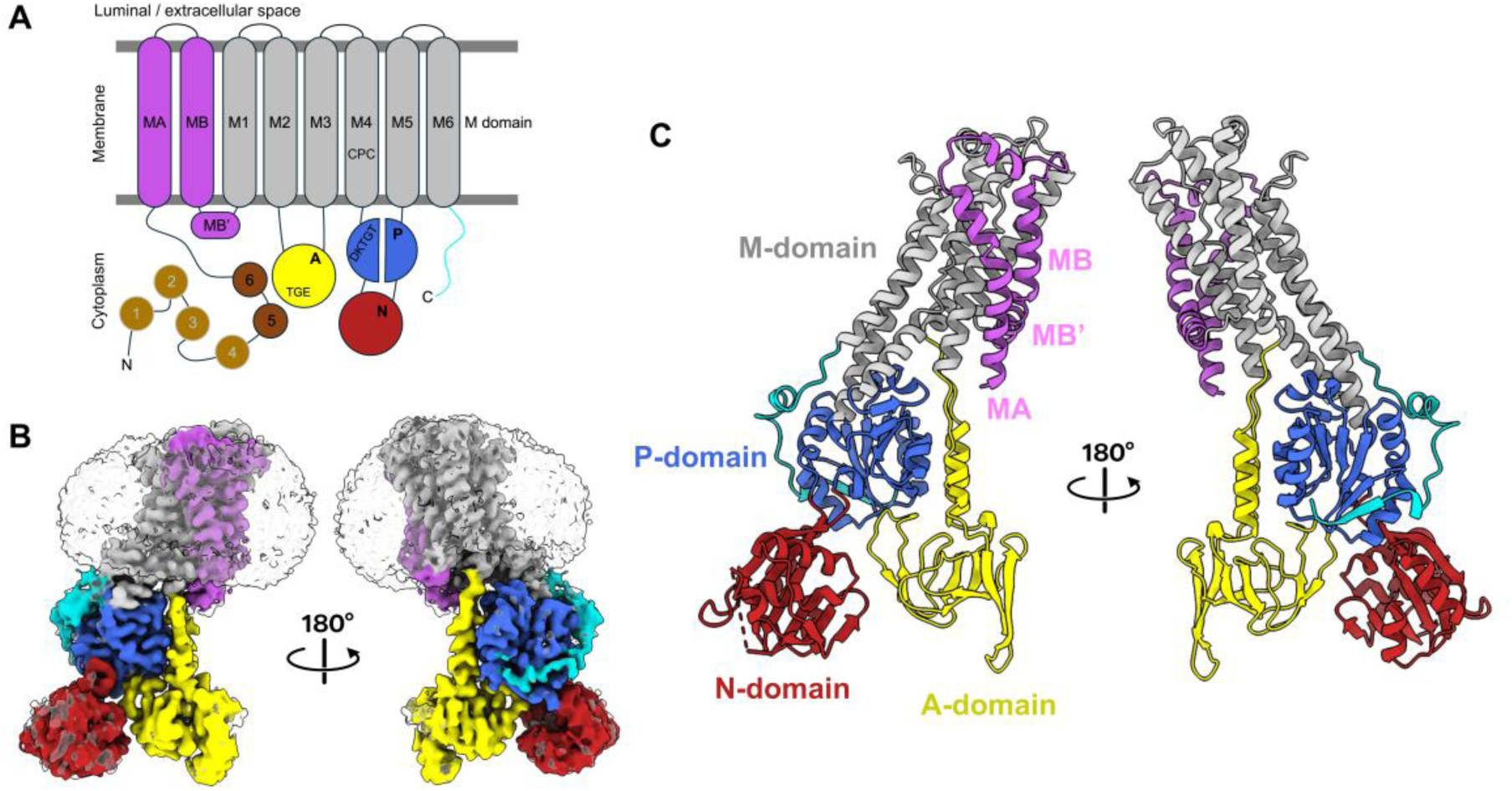
Topology, domain organization and overall structure of human ATP7B. A) Membrane topology diagram with MBDs (metal binding domains) in shades of brown, M (transmembrane) domain in silver with helices MA, MB and MB’ in purple, N (nucleotide binding) domain in red, P (phosphorylation) domain in blue, A (actuator) domain in yellow and the C-terminal tail in cyan. Conserved motifs depicted in one-letter code. Colour scheme consistent throughout all figures of this manuscript. B) Cryo-EM density map of human ATP7B_Δ_ (ΔMBD1–4, Δaa 57–481) in MSP1E3D1 nanodisc (transparent) at sigma 5.2, in 180 degrees offset bilateral views. C) Structural model of ATP7B_Δ_ with transmembrane domain, cytoplasmic N-, P- and A-domains, the C-terminal tail, and one unstructured loop 1117–1141 (N domain) in 180 degrees offset bilateral views.

Additionally, human ATP7B has long terminal tails. The C-terminal tail is mostly disordered, except for a segment wrapping around the P-domain^24^, while the N-terminal tail contains six metal-binding domains (MBDs) that are separated by linker regions. The MBDs have a ferredoxin fold like the copper chaperone ATOX1 and are thought to modulate ATP7B activity by fine-tuning copper sensing^26,27^. While MBD1–4 act as an autoinhibitory element^24,28^, the MBD5-6 segment has been speculated to act as a rigid unit regulating the E1-E2 transition^29,30^. Observing MBDs via cryo-EM is challenging due to their “pearls on a string” flexible nature; observations are limited to low-quality densities for MBDs close to the core pump^24–26^, and MBD organization is still disputed^26^. Furthermore, Cu(I) is bound by chaperones or copper-chelating compounds in the cytoplasm, but whether MBDs, with CXXC copper-binding motifs and copper-chaperone-like folds, are involved in copper delivery is still debated. Clarifying such interactions is essential for understanding ATP7B autoregulation and copper delivery mechanisms, and how these are affected by disease-causing mutations^31^.

In P-type ATPases, lipid-dependent activity and regulation are well-established^32–35^. ATPase activity of *Legionella pneumophila* CopA was previously reported to be dependent on the presence of *E. coli* total lipid extract^23^ , and a homologous *Sphaerobacter thermophilus* CopB was later shown to be activated by soybean lipids^36^. LpCopA ATPase activity is specifically stimulated by the anionic lipids cardiolipin and phosphatidylglycerol, and molecular dynamics simulations highlight MA and MB helices as main interaction sites for these lipids^37^. Charge-mediated interactions were clustered around the MB’ platform, and the positively charged residues within the region are also present in human ATP7B^37^. Notably, structures of prokaryotic copper pumps have been obtained in lipid-containing environments^21–23^. In contrast, available eukaryotic structures have so far been determined from detergent-solubilized preparations^24–26^, although biochemical evidence indicates that lipids modulate ATP7B activity^26,37^.

Here, we present a cryo-EM structure of human ATP7B in a lipid nanodisc, determined in the copper-free E2-P_i_ catalytic state with the core structure, cytoplasmic domains and luminal loops of ATP7B resolved to well below 3Å. Additionally, a subset of our dataset contained lower-resolution information on MBD5-6, suggesting a copper relay mechanism in ATP7B. Our structural model led us to further produce and analyze two WD-causing variants, R969Q and W939C, located at the luminal/extracellular side of ATP7B. Although they are far from the ion and nucleotide binding sites, they are both classified as pathogenic mutations. Using a combination of functional assays, lipid-interaction focused molecular dynamics simulations and sequence analysis, we propose different WD mechanisms for the two pathogenic variants.

## Results

### Structure determination of human ATP7B

Human ATP7B (UniProt ID: P35670-1) comprises 1,465 residues, with the N-terminal ∼643 residues (∼44%) containing six metal-binding domains (MBDs) and their linkers (Fig. 1A). Because this region is prone to proteolysis, compromising sample homogeneity, and MBD1–4 mediate autoinhibition^24^ , we generated a more stable and active construct for production in *Saccharomyces cerevisiae*. Residues 57–481 were deleted from a yeast-codon-optimized ATP7B sequence to produce human Δ(MBD1-4)-ATP7B, hereafter ATP7B_Δ_. This construct retains the N-terminal peptide required for ATP7B trafficking^38,39^ and MBD5-6, which are essential for function. A comparable design was previously used for *Xenopus tropicalis* (*Xt*) ATP7B^24^ and validated for human ATP7B both *in vitro* and *in vivo*^40^.

We next considered the importance of membrane mimetics for structural studies, and we set out to determine a cryo-EM structure of human ATP7B_Δ_ reconstituted into lipid nanodiscs with the intention of identifying lipid-interaction sites. We used the membrane scaffold protein 1E3D1 (MSP1E3D1) together with bovine liver total lipid extract to reconstitute human ATP7B_Δ_ (solubilized in *n*-dodecyl-β-D-maltoside, DDM) into a near-native lipid environment. During protein purification, the addition of adenosine 5’-diphosphate (ADP) and aluminium fluoride (AlF_x_) had a stabilizing effect, as reflected by a clear transition in nano differential scanning fluorimetry melting curves (Suppl. Fig. 2). Our final dataset entailed ∼3,000 processed cryo-EM micrographs and we obtained a 3D reconstruction based on ∼150,000 particles at an overall resolution of 2.8 Å (Fig. 1B–C, Suppl. Fig. 3, 4A–B, 5, Suppl. Table 1).

### ATP7B adopts an E2-P_i_ outward-open conformation

The overall structural organization of human ATP7B_Δ_ agrees with the expected domain architecture (Fig. 1). The membrane (M-) domain consists of the N-terminal tail preceding helices MA and MB with its kinked part MB’, followed by the helices M1–6 representing the canonical P-type ATPase M-domain core^24^. Our model closely resembles the previously determined *Xt*ATP7B structure^24^ , with an RMSD of 1.33 Å across 727 aligned residue pairs (Fig. 2A). Expectedly, no copper density was observed at the canonical CPC copper-binding motif (Fig. 2B), since we did not add any to the preparation. Well-defined densities corresponding to Mg²⁺ and AlFₓ were present adjacent to the conserved phosphorylation-site residue D1027 (Fig. 2C). The ADP ligand could not be modelled with confidence based on the obtained cryo-EM map, but a partial overlap was observed for density in the nucleotide-binding pocket and the position of the adenine moiety in a crystal structure of the ADP-Mg^2+^-bound *A. fulgidus* CopA P- and N-domains, which is indicative of sub-stoichiometric or conformationally heterogeneous ADP-binding (Suppl. Fig. 6A). The transmembrane region adopts an outward-open conformation with a broad luminal pathway that narrows toward the CPC motif and is lined by a predominantly polar surface on one side and a hydrophobic surface on the other (Fig. 2D and Suppl. Fig. 6B). Together, these features support assignment of the structure to a copper-free E2-P_i_ state.

**Figure 2:**
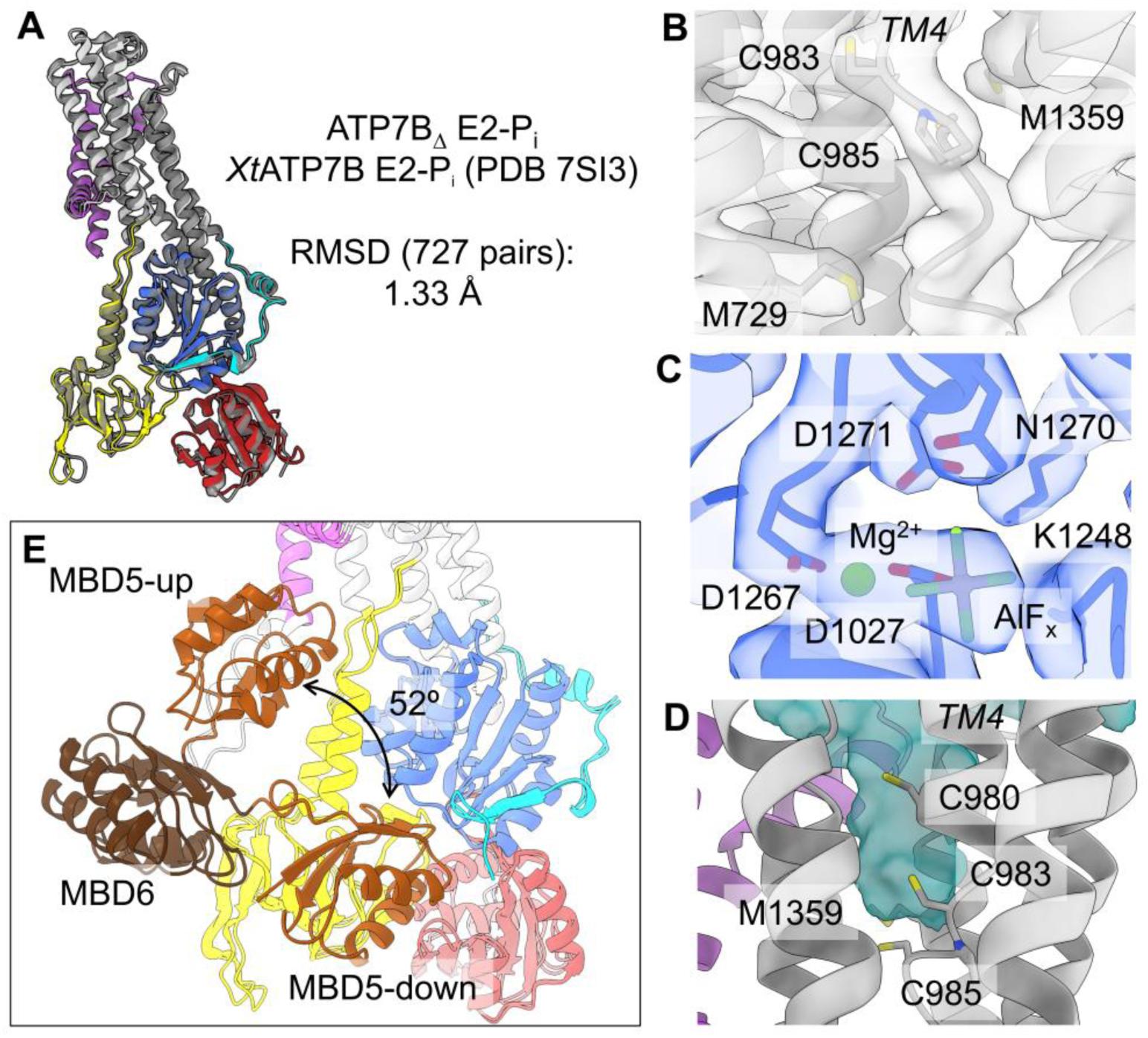
ATP7B_Δ_ in an E2-P_i_ outward-open copper-free state. A) Structure comparison of human ATP7B_Δ_ E2-P_i_ (this work) and *Xt*ATP7B E2-P_i_ (in grey, PDB ID 7SI3) with an RMSD of 1.33 Å (across all 727 pairs). B) The ion-binding site in the M-domain around the CPC motif in M4 does not contain any ions, model and map (sigma 9.5) with highlighted residues involved in copper coordination. M3 was removed for visibility. C) Map and model around the catalytic phosphorylation site at D1027 (sigma 9.5), part of the DKTGT motif. Clear densities for Mg^2+^ and AlF_x_ are present. D) Outward open exit pathway from the CPC motif, calculated with the ChimeraX 1.11 Find Cavities tool (inner probe 1.1 Å, outer probe 4.00 Å, grid spacing 0.5)^41^. E) Overlay of MBD5-down and novel MBD5-up states, depicting the 52° upward movement of MBD5 between states. Angle calculated between the Cα atom of the first cysteine in the CXXC motifs within MBD6 (C575) and MBD5 (C499) in the up and down states.

Alignment of helices MA-MB-M1 between our obtained structure and an E1 human ATP7B structure reveals a wider space between MB’ and MA in the E1 state (Suppl. Fig. 6C). This could be important for the accommodation of a copper-delivering chaperone or MBD under the MB’ helix in the cytoplasmic-open E1 state. While in previously determined structures, MBD5 likely occupies this position, we do not see this interaction here for the E2-P_i_ state^25,26^. Nevertheless, extensive 3D particle sorting of our dataset revealed a subset of particles with densities corresponding to MBD5 and MBD6 in two distinct configurations and positioning supported by clear MBD5-MBD6 and MBD6-MA linker assignments (Fig. 2E, Suppl. Fig. 3, 4C-F). In an MBD5-down configuration (Suppl. Fig. 7A), MBD6 is localized on the A-domain and MBD5 is positioned between the A- and P-domains with contacts to the C-terminal tail, as also observed for *Xt*ATP7B^24^ (Suppl. Fig. 7C). Because the A-domain provides structural support for MBD5 to reach both the P-domain and the C-terminal tail, this MBD configuration is clearly coupled to the E2 conformation. In a novel MBD5-up configuration (Suppl. Fig. 7B) MBD5 shifts from the A-/P-domain cleft upward by about 52° toward the MB’ platform (Fig. 2E), while MBD6 occupies a similar position in both states. The spatial constraints of the E2-P_i_ conformation prevent MBD5 from fully reaching the copper-entry cavity. Instead, MBD5 appears to rest on the MBD6-MA linker, resembling a state primed for docking into the M-domain (Suppl. Fig. 7B,D). Together, the previously reported MBD5-down state and the MBD5-up state described here reveal conformational heterogeneity within the ATP7B E2-P_i_ state and suggest a possible pathway by which MBD5 becomes positioned for copper delivery to the M-domain in subsequent E1 states.

### Wilson disease mutation R969Q disrupts a functionally important salt bridge

The increased resolution in the periphery of our map obtained from nanodisc-reconstituted ATP7B, especially around the long luminal loops, allowed us to build a complete structural model for this region. Two WD mutants locate here: W939C (dbSNP^9^ ID: *rs1057517310*) and R969Q (dbSNP^9^ ID: *rs121907996*) (Fig. 3A, Suppl. Table 2). The latter mutant has previously been shown to cause reduced copper transport activity and hyperphosphorylated ATP7B but exhibited WT-like activity in a tyrosinase assay^42,43^. Taken together with its limited conservation, its benign AlphaMissense prediction (Suppl. Fig. 8D-E) and the residue location at the periphery of the pump, the mechanism by which R969Q influences ATP7B in WD was elusive. Hence, we proceeded to produce and purify the R969Q variant and compared it to WT-ATP7B_Δ_ and the catalytically inactive D1027N and WD variant H1069Q in an ATPase assay. Importantly, we found it to be completely inactive (Fig. 3A-B, Suppl. Fig. 8A-C).

**Figure 3:**
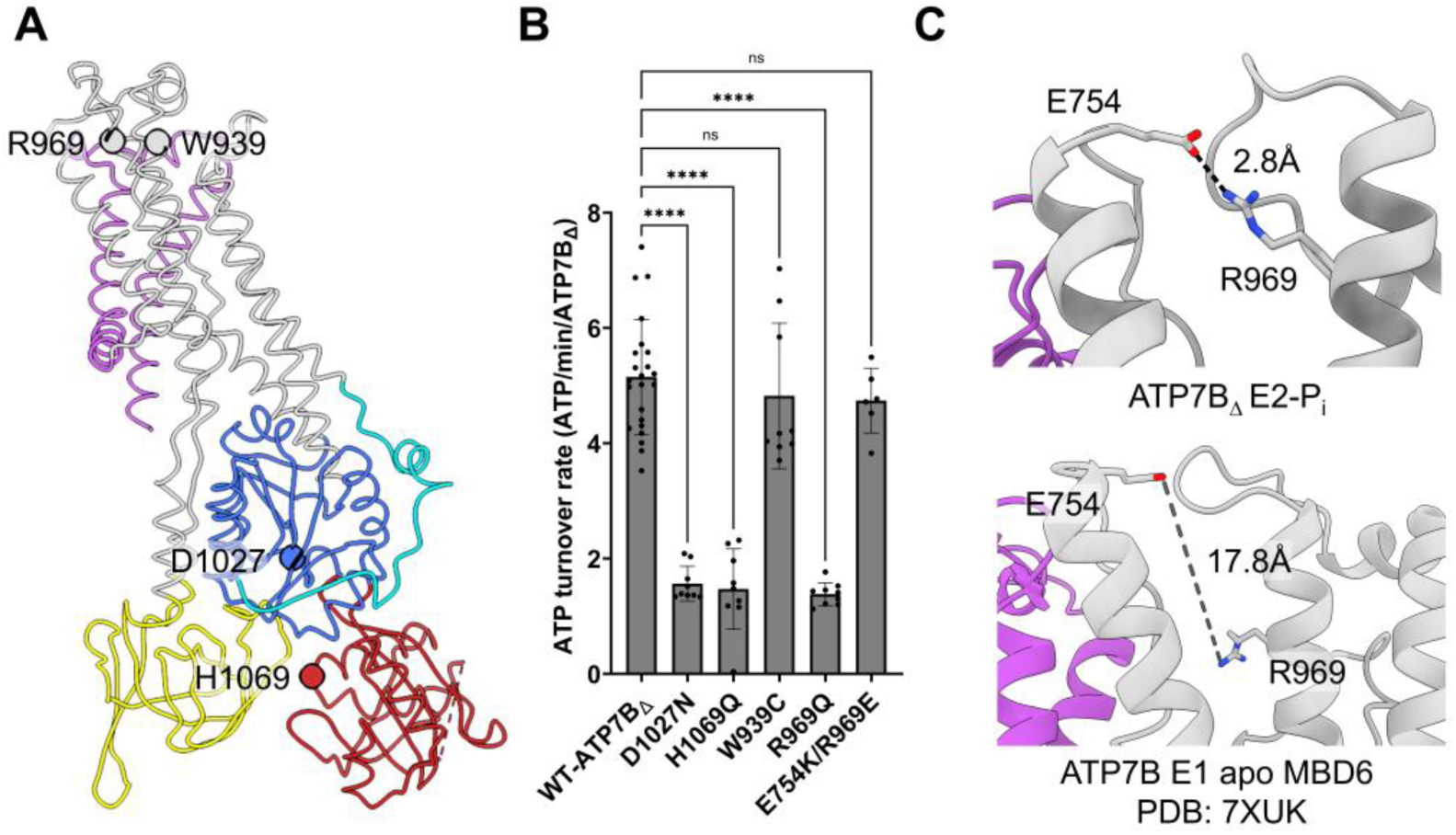
Mutations in ATP7B cause Wilson disease. A) Localization of ATP7B residues marked by Cα spheres. Highlighted are the investigated WD mutants W939C and R969Q, as well as the most common WD variant in Europe H1069Q and the catalytically inactive ATP7B variant D1027N. B) ATP turnover rates of WT, catalytically inactive (D1027N), three WD mutations of ATP7B and double mutant E754K/R969E. Dots represent individual measurements, one-way ANOVA comparing the mean of biological replicates was done followed by Dunnett’s multiple comparisons test (****adjusted p-value <0.0001). At least three biological replicates/variant, except E754K/R969E, where two biological replicates were measured. C) Close-up view of the salt bridge between R969 and E754 present in ATP7B in an E2-P_i_ conformation (our study). The same residues are 17.8Å apart in ATP7B in an E1 apo conformation (PDB:7XUK).

In our E2-P_i_ structure, R969 (M4) forms a previously unrecognized salt bridge with E754 on opposing helix M1 (Fig. 3C, Suppl. Fig. 9A, D). This interaction is disrupted in the E1 conformation of ATP7B. Remarkably, R969 is only conserved in higher eukaryotes (Suppl. Fig. 8D, 9B), but an inverted salt bridge (E583 and K373) is found in the rice homologue HMA4, i.e. conserving the salt bridge across species (Suppl. Fig. 8D, 9C). Introduction of the inverted salt bridge into WT-ATP7B_Δ_ (E754K/R969E) conserved ATPase activity (Fig. 3B, Suppl. Fig. 8A), further underlining the importance of this interaction for a functional cycle. From our structural observations, ATPase assays and previous analyses, especially the hyperphosphorylation study, we conclude that the R969Q mutation prohibits the formation of the salt bridge and likely stalls the E1P to E2P transition of ATP7B and renders it inactive.

### WD mutation W939C disrupts a crucial lipid interaction site

The second examined mutation, W939C, exhibited ATPase activity comparable to that of WT-ATP7B_Δ_ (Fig. 3B), but protein yields were greatly reduced when compared to the other ATP7B variants (Suppl. Fig. 8A-C). In the structure of ATP7B_Δ_ in an MSP1E3D1 nanodisc, a flat, elongated lipid-like density is observed near W939 (Fig. 4A).

**Figure 4:**
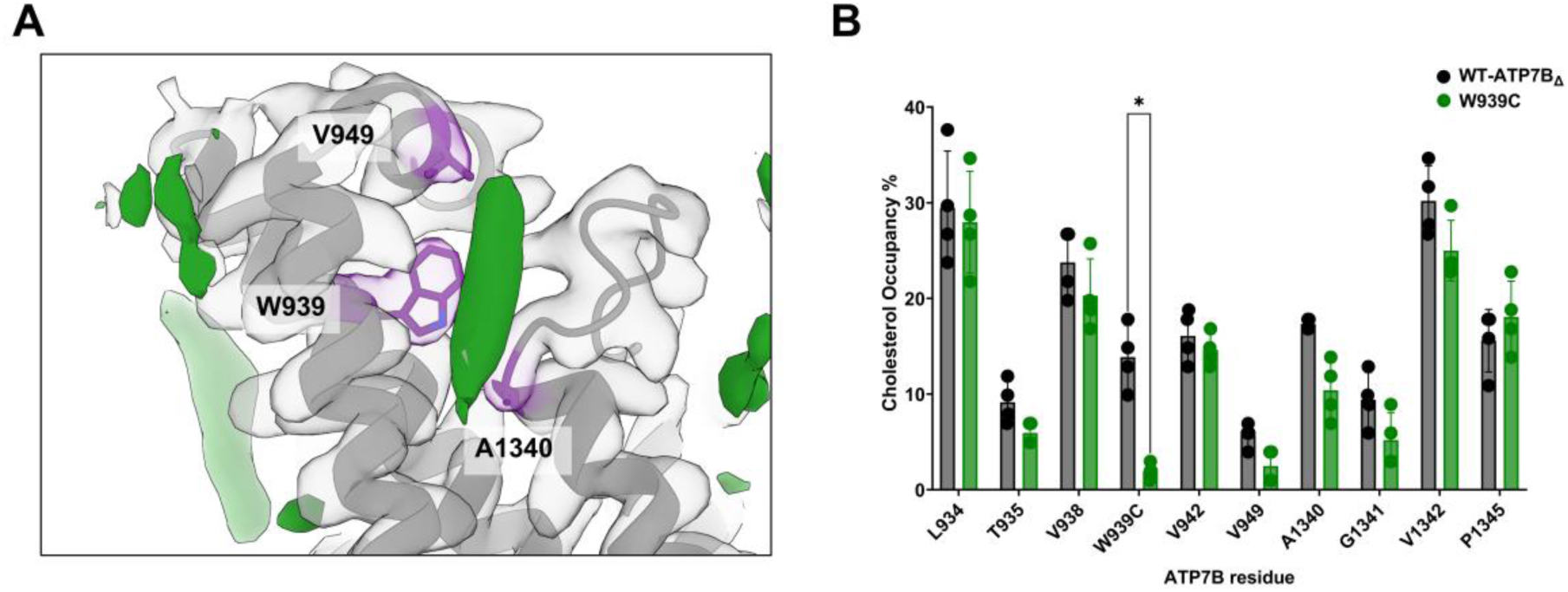
Analysis of a unique cholesterol-binding site found in ATP7B. A) Cryo-EM map and model of human ATP7B_Δ_ around W939, with a flat lipid-like density (green) embedded between M3 and M5. B) Comparison of mean cholesterol occupancy % between WT and W939C ATP7B_Δ_ MD-simulations for residues surrounding the lipid density observed near W939 in our cryo-EM map (A). Data from four independent 10 µs simulations with WT-ATP7B_Δ_ and W939C-ATP7B_Δ_ inputs. Occupancy is presented as mean ± SD. Statistics: Benjamini and Hochberg false discovery rate (FDR) of 5% was applied after Welch’s t tests. A significant difference (*) was observed only for W939C (q=0.0427), while V949 and A1340 had q-values below 0.1.

We used the Cholesterol Binding Domain (CBD) Finder tool^44^ to identify putative cholesterol binding motifs in ATP7B (Suppl. Fig. 10). All identified sites in the M region of ATP7B are in proximity of flat lipid-like densities in our cryo-EM map. However, the lipid-like density next to W939 was not identified in this analysis. Since a comparable density is present in the cryo-EM maps of detergent/CHS-solubilized *Xt*ATP7B, inactive human ATP7B and *Os*HMA4 (Suppl. Fig. 11), and since W939 is highly conserved (Suppl. Fig. 8D), we propose it could be a unique sterol-binding site of ATP7B homologues. To test this model, we performed coarse-grained molecular dynamics simulations (CG-MD) of WT-ATP7B_Δ_ and analyzed lipid occupancy around ATP7B, especially in the vicinity of W939. We further included W939C-ATP7B_Δ_ in this analysis to assess whether this mutation (predicted to be highly pathogenic by AlphaMissense (Suppl. Fig. 8E)) influenced lipid occupancy.

We performed CG-MD simulations of ATP7B in a lipid bilayer (40% 1,2-dioleoyl-sn-glycero-3-phosphocholine (DOPC), 40% 1-palmitoyl-2-oleoyl-glycero-3-phosphocholine (POPC), 20% cholesterol (CHOL)) with our structural model of WT-ATP7B_Δ_ as input. Lipid interactions across ATP7B were analyzed via the LipIDens protocol^45^ between four 10 µs simulations (see workflow in Suppl. Fig. 12A). The overall lipid occupancies matched the residues in hydrophobic patches of the transmembrane region (Suppl. Fig. 12B). Cholesterol occupancy was high (28-34%) for several of the conserved residues in the predicted cholesterol binding motifs (e.g. F654, Y715, F972; Suppl. Fig. 10, 12). Apart from the predicted cholesterol-binding motifs, cholesterol occupancy was increased for residues around W939 (Fig. 4B), all of which are close to the cholesterol-like density observed in our cryoEM map. To test whether the W939C mutation disrupts this potential cholesterol interaction, four simulations were performed using the W939C-ATP7B_Δ_ mutant as an input, utilizing the same protocol as for WT-ATP7B_Δ_. Differences in lipid occupancy were the highest in the case of residue 939. At this position, cholesterol occupancy decreased from 14% (WT, W939) to 2% (mutant, 939C). Targeted statistical analysis of lipid occupancy at residues in the proximity of the cholesterol-like lipid density near W939 showed that only this difference was significant (Fig. 4B). Nevertheless, the trend of decreased cholesterol occupancy was also observed for other residues near the lipid density at this site, namely T935, V938, V949 on M3 and A1340, G1341, V1342 on M5 (Fig. 4B). Notably, no such trend was observed for POPC or DOPC occupancy of the same set of residues (Suppl. Fig. 12C). Overall, changes in lipid occupancy were minimal between the WT and W939C simulations apart from the cholesterol occupancy around residue 939, described here. Our CG-MD simulations therefore show that the WD mutation W939C influences a crucial ATP7B-cholesterol interaction. Overall, our nanodisc-based approach revealed a previously overlooked cholesterol-binding site near W939 and supports the view that a lipidic environment is important for ATP7B stability and function.

## Discussion

WD is typically a manageable disease, but it still requires lifelong daily treatment. Although numerous pathogenic ATP7B variants have been identified, many aspects of ATP7B function remain understudied and the mechanisms by which disease-associated substitutions influence ATP7B are incompletely understood. Here, our cryo-EM structure of human ATP7B in an E2-P_i_ conformation sheds new light on a possible copper-delivery pathway and, together with *in vitro* activity assays and *in silico* lipid analysis, provides mechanistic insights into the disease-associated variants R969Q and W939C.

### H1069Q and possible modulatory nucleotide binding

In our sample preparation, the addition of ADP and AlFₓ in the absence of supplemented or endogenous copper led to ATP7B stabilized in an outward-open, copper-free E2-P_i_-like conformation. Because catalytic phosphorylation of ATP7B is copper-dependent^46^, we speculate that the absence of copper restricted conversion to the tightly nucleotide-engaged E1 state. The partial density we observe adjacent to H1069 (Suppl. Fig. 5A) could instead reflect a weak or modulatory nucleotide-binding mode analogous to the E2 state described for SERCA^47,48^. Copper binding could therefore facilitate conversion from a weakly nucleotide-bound E2-P_i_-like state to an E1 state competent for phosphoryl transfer. An H1069Q substitution likely disrupts this coupling since it reduces nucleotide affinity and destabilizes the N-domain^49–52^.

### Regulation at the A-, N-domain, C-terminal tail and MBD5 junction

Additionally, copper binding induces rearrangements of the N-terminal regulatory region^31,53^. Particularly, MBD5 has been shown to move from the space between the A- and P-domains (E2-P_i_) ^24^ to under the MB’ platform (E1) in a state-dependent manner^25,26^. On the other hand, a role of MBD6 as a build-on, static module of the A-domain seems plausible^26^. Applying extensive 3D particle sorting to our data, we were able to extract two cryo-EM maps reflecting a rather static MBD6 on the A domain and distinct configurations of MBD5, which we term MBD5-down and MBD5-up (Fig. 2E, Suppl. Fig. 7). This separation came at the cost of high-resolution detail because of the intrinsic flexibility of the MBDs and the reduced particle numbers. The previously observed MBD5-down state seems to be induced by interactions with the A- and P-domains and the C-terminal tail^25^. The C-terminal tail has been proposed to support regulatory functions associated with intracellular compartmentalization and organelle-specific trafficking^24,54^. The short MBD5-MBD6 linker, acting as an elbow, restricts MBD5 from reaching towards MB’ or to interact with the C-terminal tail. The upward movement of MBD5 in our novel MBD5-up configuration is limited by the E2-P_i_ conformation, and it does not reach MB’, as in E1 conformations. Instead, it settles on top of the MBD6-MA linker. Interestingly, this linker contains a cluster of WD mutations, indicating its importance in ATP7B function^55^. The following E2 to E1 transition requires the separation of the A- and P-domains, which moves the A-domain/MBD6 segment and associated helices by ∼3 helical turns upwards to provide space under MB’ for MBD5 docking (Suppl. Fig. 6C, 9D)^25^. We speculate that especially MBD5, with MBD6 as an anchor, facilitates and stimulates ATP7B conformational changes, and that the ‘MBD5-down’ configuration is not autoinhibitory, as previously discussed, but allows for regulatory contacts with the C-terminal tail, consistent with several truncation / mutational studies (Suppl. Table 3).

### A crucial salt bridge stabilizes the E2-P_i_ conformation

While earlier conflicting data on the functionality of the R969Q mutation existed, it was not clear how this mutation, which is localized far from the central ion and nucleotide binding sites, could influence ATP7B activity. R969Q was shown to have wild-type-like activity in a tyrosinase assay^43^ and was predicted to be a benign mutation by AlphaMissense, although it leads to decreased ^64^Cu-uptake in membrane vesicles and hyperphosphorylation^42^. Moreover, mass-spectrometry-based analyses identified patients with Wilson disease carrying R969Q, despite wild-type-like ATP7B protein abundance^56^. Additionally, WT-like intracellular localization has been reported for this mutant^42,57^. Although less common than leading disease-associated variants such as H1069Q and R778L, R969Q is frequently identified in patients with WD, and despite the clear connection with WD, it is sometimes identified as normal/WT-like in clinical screens, underscoring the importance of understanding its molecular mechanism^58^.

We found R969Q to be inactive *in vitro*, and through careful structural comparisons, we identified the crucial role of R969 (M4) in the formation of a salt bridge with E754 (M1). This salt bridge seems to stabilize the E2-P_i_ state of ATP7B and is most likely needed for the E1P-to-E2P transition, as its disruption abolishes ATP turnover, while a charge inversion maintains full activity. Interestingly, there seems to be a correlation between the presence of the salt bridge and a higher number of MBDs (at least three) per copper pump. Together, these observations reconcile the previously contradictory findings on R969Q and support a novel WD mechanism acting on ATP7B function. During ATP7B turnover, the formation and breaking of this salt bridge are coupled to the movements of MBD5, the elbow linker between MBD5-6 and MBD6 sitting static on the A domain. The separation of A- and P-domains and the docking of MBD5 below MB’, will push MB and M1 upwards and break the salt bridge (Suppl. Fig. 9D). These large structural changes of the core pump, MBDs and reiterative reformation of the salt bridge are high-energy expenses. But ATP7B is an extremely slow pump (turnover of <10 ATP/min/protein) compared to ultra-fast ion pumps like SERCA and NKA (up to 9000 ATP/min/protein)^24,59,60^. Additionally, copper is a scarce (1–2 mg/kg body weight) but essential trace element that needs to be tightly regulated due to cell toxicity^61,62^. Hence, it is always complexed by copper chaperones or MBDs. This proposed coupling mechanism directly links copper availability to ATP7B turnover, limiting futile pump activity and energy expenses. Accordingly, therapeutic correction of this mutation would likely require either gene editing/replacement or a drug that reinforces E1-E2 cycling.

### Identification of a non-canonical stabilizing cholesterol binding site

Like for R969Q, the pathogenic mechanism of W939C is difficult to infer from sequence or position alone as it is far from obvious functional sites. To our knowledge, there is little prior experimental data available on this variant. W939C exhibits WT-like ATPase activity but we noted a much lower protein yield, compared to the other purified mutants (Suppl. Fig. 8A–C). W939 is highly conserved (Suppl. Fig. 8D) across species, and a lipid-like density close to W939 is observed in several cryo-EM maps of ATP7B and related copper pumps (Suppl. Fig. 11). Our CG-MD simulations show that W939C reduces cholesterol occupancy in the cleft between M3 and M5 formed around that residue, and we propose that the binding of cholesterol or substitute sterol (ergosterol or CHS) is crucial. The sterol likely acts as a folding chaperone, stabilizing ATP7B and allowing for proper trafficking and preventing protein degradation. Such a mechanism would make the cleft between M3 and M5 an interesting target for a small-molecule binder, reverting the destabilization of W939C and other WD mutants (e.g. T935M, G943D/R/S, and G1341D/R/V) in this area (see Suppl. Table 2 for details on mutants).

Indeed, G943D/S mutants were previously shown to be active, albeit expressed at lower levels than the WT protein, and the G943S mutant was unable to traffic properly upon copper treatment^63–65^. Normal activity and localization have been reported for the T935M mutant; however, its expression level was lower than that of the WT protein^63,66^. The G1341V mutant also exhibited severely reduced protein abundance and copper export that was partially rescued by pharmacological chaperones (4-PBA and curcumin) restoring protein levels^67^. Together, these observations suggest a common mechanism underlying these mutations, based on a defect in protein stability rather than catalysis, in which loss of cholesterol engagement in the M3-M5 cleft may destabilize ATP7B and impair its folding and/or trafficking. Our data pinpoint the cholesterol-binding cleft near W939 as a potential druggable site for stabilizing ATP7B and rescuing W939C and related WD variants, opening a previously unrecognized therapeutic avenue (Suppl. Fig. 13).

### Potential strategies for correction of WD variants

Collectively, the presented and previous findings suggest that ATP7B variants will likely require mechanism-specific correction strategies. Variants that affect catalytic cycling (e.g. R969Q) may require restoration of productive conformational transitions, whereas destabilizing variants (e.g. W939C) may be more amenable to pharmacological stabilization of vulnerable structural elements such as lipid- or sterol-associated sites. A ligand occupying the lipophilic M3-M5 pocket could in principle compensate for reduced sterol binding and act as a pharmacological chaperone, analogous to type I or III CFTR correctors^68^. The partial rescue of H1069Q by reduced temperature, chemical chaperones and suppression of mutant-protein degradation further supports stabilization-based correction as a feasible therapeutic principle for ATP7B ^14,16,67,69,70^. Together, in addition to the two novel WD mechanisms centred on W939 and R969 residues, our study advances the general understanding of the ATP7B transport mechanism. Specifically, it provides the first structure of human ATP7B in an E2 conformation, capturing the opposite side of the Post-Albers cycle.

## Conclusion

Our cryo-EM structure of human ATP7B in an E2-P_i_ conformation determined at 2.8 Å overall resolution in a liver lipid nanodisc complements the earlier structures of inactive ATP7B in E1 conformations^25^ and enables detailed analysis of the E1-E2 transitions of ATP7B. Our structure reconciles previous disagreements regarding the roles and localization of MBD5 and MBD6 relative to the core pump. In particular, we identify a previously undescribed MBD5-up configuration, in which MBD5 appears primed for docking beneath MB’ prior to the E2-E1 transition. The same upward movement of MBD5 is crucial in understanding the inactivating mechanism of WD variant R969Q affecting a salt bridge that stabilizes E2 states of ATP7B. Finally, our data highlight the importance of structural lipids for proper ATP7B function and suggest that the identified lipid-binding groove around WD variant W939C may provide a favourable site for small-molecule stabilization of disease-associated ATP7B variants.

## Limitations

One limitation of our study is that our presented ATPase assays are independent of added copper substrate. Hence, the values we present represent a basal ATPase activity. The slow turnover we observe for human ATP7B_Δ_ (∼6 ATP/min/protein) matches well with the one measured by Bitter *et al*. for a similar Δ(1-4MBD)-ATP7B from *Xenopus tropicalis* (7.8 ± 0.5 ATP/min/protein)^24^ , and our measured activity is clearly inhibited via the addition of 1 mM P-type ATPase inhibitor BeF_x_ (Suppl. Fig. 8A). Ours and other trials to add copper have resulted in inhomogeneous samples and/or heavy precipitation^25^, and to date, there is no report of direct, copper-dependent stimulation of ATP7B ATPase activity *in vitro*. As the mutations we evaluate in our ATPase assay are all far away from the central ion-binding sites around the CPC motif, we are confident that the clear trends in ATPase activity we report (WT-ATP7B_Δ_-like activity or DN mutant-like inactivity) will persist in different setups.

Another limitation of our study is that we cannot discriminate between ergosterol, cholesterol and cholesterol hemisuccinate (CHS) in many of our experiments. The lipid densities we see in our cryo-EM map can be either of the three sterol lipids, as ergosterol could be co-purified from our *Saccharomyces cerevisiae* expression system, CHS is added during ATP7B solubilization and cholesterol is present in the bovine total liver extract used during nanodisc reconstitution. During ATPase assays, both co-purified ergosterol and CHS could be present. The only sterol present in the CG-MD simulations was cholesterol. We, however, remain confident in our cholesterol-related conclusions, as cholesterol is the native sterol ATP7B is exposed to, and our cholesterol-binding site analysis and the cholesterol occupancies from the CG-MD simulations match the cholesterol-like densities in our cryo-EM map.

## 1 Methods

### 1.1 DNA constructs

The original pYeDP60 vector backbone for galactose-induced protein production in yeast was a gift from Guillaume Lenoir’s lab. The human ATP7B gene, encoding a sequence with UniProt ID P35670-1 containing a residue 57–481 deletion (ΔMBD1–4) and polymorphisms K832R, R952K, and V1140A, was obtained from GenScript. The sequence was codon-optimized for *Saccharomyces cerevisiae,* together with a C-terminal eGFP tag separated from ATP7B by the TEV protease cleavage site. The Δ(MBD1-4)-ATP7B-eGFP construct flanked by EcoRI and SacI sites in the pUC57 subcloning vector was cut with the corresponding enzymes and ligated into the pYeDP60 vector. The construct to produce membrane scaffold protein 1E3D1, pMSP1E3D1, was a gift from Stephen Sligar^71^ (Addgene plasmid #20066).

Site-directed mutagenesis was performed on the pUC57 vector to obtain H1069Q and D1027N mutants of ATP7B using the QuikChange Lightning kit (Agilent Technologies, #210519). Subsequently, the mutants were cloned into the pYeDP60 vector as described above for the wild type. The same kit was used to generate W939C and R969Q mutants, but directly using the pYeDP60-Δ(MBD1-4)-ATP7B-eGFP plasmid. To generate the double mutant E754K/R969E, GeneArt™ Gibson Assembly HiFi Cloning Kit (Thermo Fisher Scientific, #A46624) and the same pYeDP60 vector was used as a backbone and template for the mutagenesis. All generated constructs were validated via whole plasmid sequencing. The primers used are listed in Suppl. Table 4.

### 1.2 Production and purification of MSP1E3D1

MSP1E3D1 was produced in the BL21(DE3) *Escherichia coli* strain (Thermo Scientific, #EC0114). Once the cell culture, grown at 37 °C and 120 rpm in TB medium (1.2% w/v tryptone, 2.4% w/v yeast extract, 0.5% v/v glycerol, 17 mM KH_2_PO_4_, and 72 mM K_2_HPO_4_) with 50 μg/mL kanamycin, reached an OD600 of 0.7, protein production was induced by 0.2 mM IPTG following a 10 min incubation in an ice bath. Cells were harvested after 16 h cultivation at 18 °C and 120 rpm and stored as pellets at - 70 °C. Cells were lysed by sonication in a buffer containing 50 mM Tris-HCl pH 8.0, 500 mM NaCl, 20 mM imidazole, 1 mM PMSF, 3 mM MgCl_2_, and 10 μg/mL DNase I. The lysate was clarified by centrifugation at 20,000×g for 30 min at 4 °C. The supernatant loaded under gravity flow onto a Ni Sepharose 6 Fast Flow resin (GE Healthcare, #17-5318-02) equilibrated in lysis buffer and used at a ratio of 1.5 mL of beads per 10 g of wet weight of the input cell pellet. After binding, the resin was washed using 20 CV of buffer containing 50 mM Tris-HCl pH 8.0, 500 mM NaCl, and 20 mM imidazole. The same buffer but containing 500 mM imidazole was used for elution, and pooled protein-containing fractions were dialysed against buffer containing 50 mM Tris-HCl pH 8.0, 150 mM NaCl, 10 mM imidazole, and 1 mM DTT. His-tag was cleaved from MSP1E3D1 using His-tagged HyperTEV60 protease added during dialysis at a 1:20 weight ratio. The tag and protease were separated from pure MSP1E3D1 by reverse IMAC procedure using the same regenerated resin and dialysis buffer. The protein was concentrated to 20–30 mg/mL and stored at -80 °C. HyperTEV60 protease was produced and purified as described in ^72^, and its DNA construct was a gift from Magnus Kjærgaard.

### 1.3 Production of Δ(1-4MBD)-ATP7B-EGFP and yeast membrane preparation

The yeast strain used in this study, *Saccharomyces cerevisiae* W303.1b/Δ*pep4* (*α*, *leu2-3*, *trp1-1*, *his3, pep4*Δ*:loxP, ura3-1*, *ade2-1*, *can*^r^, *cir*^+^), was a gift from Rosa Lopez Marquez. Yeast cells were transformed using the LiAc/SS-DNA/PEG procedure^73^. Yeast pre-cultures were grown in SD-URA medium comprising 1× yeast nitrogen base (US Biological, #Y2025), 1× dropout medium (CSM-Ura w/ Ade40; Sunrise Science, #1103-100), and 2% (w/v) D-(+)-glucose. For protein production, the culture was upscaled in a rich medium and induced with galactose in two steps^74^. Cells were harvested at 4,000×g for 10 min at 18 °C, washed in ice-cold water, and resuspended in 2 mL of TEKS buffer (50 mM Tris-HCl pH 7.5, 100 mM KCl, 1 mM EDTA, and 600 mM D-sorbitol) per 1 g of cell pellet wet weight. After a 15 min incubation on ice, the cells were spun at 4,000×g for 10 min at 4 °C, flash-frozen in liquid nitrogen, and stored at -70 °C. 30–40 g of wet cell weight was obtained per 1 L of culture.

Membranes from the harvested cells were prepared using bead beater homogenizers with 0.5 mm glass beads. Cells were resuspended in TES buffer (50 mM Tris-HCl pH 7.5, 1 mM EDTA, and 600 mM D-sorbitol) with 2 mM PMSF and 2 µg/mL of each leupeptin, pepstatin A, and chymostatin. After 8 cycles of 1 min bead-beating and 1 min cooling, the homogenate was supplemented under stirring with 3 µL of 10 M NaOH per 1 g of starting cell weight, and subsequently clarified at 4,000×g for 10 min at 4 °C. Membranes were harvested from the supernatant by pelleting at 177,000×g for 1.5 h at 4 °C. The resulting pellet was resuspended in TES buffer with protease inhibitors at a ratio of 0.2 mL per 1 g of starting cell weight and stored at -70 °C after flash-freezing in liquid nitrogen. Total protein concentration in the membranes was determined using a BCA assay kit (Thermo Scientific, #23225) as recommended by the manufacturer.

### 1.4 Purification of ATP7B_Δ_

ATP7B_Δ_ was purified in *n*-dodecyl-β-D-maltoside (DDM) with cholesteryl hemisuccinate (CHS) via anti-GFP affinity chromatography with proteolytic elution. Briefly, Δ(1-4MBD)-ATP7B-eGFP construct-containing membranes at a concentration of total protein of 5 mg/mL were stirred at 8 °C for 1 h in purification buffer (50 mM MOPS-Tris pH 7.0, 150 mM NaCl, 5 mM MgCl_2_, and 0.5 mM TCEP) with 20% w/v glycerol, 0.5% w/v DDM, 0.1% w/v CHS, and protease inhibitors (1 mM PMSF and 1 μg/mL of each leupeptin, pepstatin A, and chymostatin). The mixture was clarified at 177,000×g for 1 h at 4 °C, and 15 µL of anti-GFP resin (prepared in-house as described in^75^) per 1 mL of obtained supernatant was added and incubated under agitation for 1 h at 8 °C. Resin was packed into a gravity column and washed under flow with 1×5 column volumes (CV) of wash buffer (same as solubilization buffer but with 0.05/0.01% w/v DDM/CHS) and 2×5CV washes without protease inhibitors. To elute ATP7B_Δ_, the eGFP tag was cleaved by HyperTEV60 protease added in a ratio of 70 µg per 1 mL of anti-GFP resin. Resin was incubated with the protease for 16 h under agitation at 8 °C and the protein was eluted under gravity flow using 3×1CV of wash buffer without inhibitors. After concentrating to ∼5 mg/mL using 100 kDa MWCO filter (Millipore, #UFC810008), the eluate was injected onto a Superose 6 10/300 column (Cytiva, #29091596) with purification buffer used as the running buffer. SEC-purified monomeric fraction was used in reconstitutions and activity assays.

### 1.5 ATP7B_Δ_ reconstitution into MSP1E3D1 nanodisc

Reconstitution was done using a SEC-purified monomeric sample (0.5–0.7 mg/mL protein) solubilized in DDM/CHS obtained as described above. Lipids from bovine total liver extract (Avanti, #181104P-100MG-D-055) were used with 750 Da as an approximate MW. Reconstitution mixture comprised an ATP7B/MSP1E3D1/lipid molar ratio of 1:8:60 in reconstitution buffer (50 mM MOPS-Tris pH 7.0, 150 mM NaCl, 5 mM MgCl_2_, and 0.5 mM TCEP) with 0.05/0.01% w/v DDM/CHS. Protein was first incubated with lipids for 15 min, then with MSP for 30 min while both incubations occurred under 15 rpm rotation at 25 °C. Detergent was depleted using Bio-Beads SM2 (Bio-Rad, #1523920) added in a ratio of 30 mg of equilibrated Bio-Beads per 1 mg of DDM for 12 h under 15 rpm rotation at 8 °C. Reconstituted sample was concentrated using a 100 kDa MWCO concentrator and refined by SEC on a 10/300 Superdex 200 Inc. column. Monomeric fraction of ATP7B-MSP1E3D1 nanodisc was collected, concentrated, and stored at -80 °C upon flash-freezing.

### 1.6 Fluorescence detection size exclusion chromatography (FSEC)

We previously reported this method and corresponding results in^76^. Briefly, DDM- and DDM/CHS-solubilized crude membranes were used for the FSEC analysis. A solubilization mixture containing 5 mg/mL total protein was prepared as described above, except that a detergent concentration of 1.25% (w/v) was used. Samples were clarified at 100,000×g for 30 min at 4 °C and 100 µg of total protein was injected in 20 µL onto a Superdex 200 Increase 5/150 GL column (Cytiva, #28990945) coupled to a Shimadzu Prominence Modular HPLC system with a UV-VIS detector SPD-20A and a fluorescence detector RF-20A. The running buffer comprised 50 mM MOPS-Tris pH 7.0, 150 mM NaCl, 5 mM MgCl₂, 0.5 mM TCEP, and 0.026% (w/v) DDM, with 0.005% (w/v) CHS included for DDM/CHS-solubilized samples. Fluorescence was detected with an excitation wavelength of 488 nm, an emission wavelength of 512 nm, high sensitivity, and 1× gain.

### 1.7 ATPase assay

ATPase activity of purified ATP7B_Δ_ samples was quantified using a malachite green (MG) assay. MG working solution was prepared under stirring and cooling on ice by combining 50 volumes of MG stock solution (3 M H_2_SO_4_, 1.22 mg/mL malachite green), 1 volume of 11% (v/v) Tween 20, and 12.5 volumes of 7.5% (w/v) ammonium molybdate tetrahydrate. Total volume of 12.5 µL of reaction buffer (15% v/v glycerol, 50 mM MOPS-Tris pH 7.0, 150 mM NaCl, 5 mM MgCl_2_, 5 mM KCl, 0.5 mM TCEP, and 0.05/0.01% w/v DDM/CHS) with 3 µM ATP7B sample was equilibrated on ice for 2 min with or without the ATP7B inhibitor BeF_x_, used in 1 mM final concentration. The reaction was started by addition of ATP to a final concentration of 3 mM. After 60 min at 37 °C, 62.5 µL of ice-cold 128 mM H_2_SO_4_ was added and the samples were incubated on ice for 10 min to stop the reaction. Following the addition of 25 µL of MG working solution and incubation for 10 min at 25 °C, the reaction development was stopped by adding 25 µL of 17% (w/v) trisodium citrate dihydrate solution. Absorbance at 620 nm of the final mixture (175 µL) was read in a SpectraMax i3x Multi-Mode Microplate Reader (Molecular Devices). In all cases, at least three biological replicates in three technical parallels were measured, except the E754K/R969E double mutant for which only two biological replicates were measured. Individual values are presented in the figures, one-way ANOVA for biological replicates (averages of the technical parallels) followed by Dunnett’s multiple comparisons test (α=0.05) was done to compare changes in ATPase activity. Data analysis and visualization were performed using GraphPad Prism version 10.6.1.

### 1.8 Nano differential scanning fluorimetry (nanoDSF)

Prometheus Panta (NanoTemper Technologies) was used for nanoDSF measurement. Thermal unfolding within the range of 20–95 °C (1 °C/min ramp) was recorded with active DLS acquisition. The DDM/CHS-solubilized WT-ATP7B_Δ_ samples were analyzed at a concentration of 0.6 mg/mL. The protein samples in apo state or treated with 0.5 mM ADP and 1 mM AlF_x_ were obtained as described above. The apo protein dataset comprises three measurements from independent purifications, whereas the ADP-AlF_x_ dataset represents a technical duplicate of the sample used for the cryo-EM analysis. The nanoDSF data was acquired and analyzed using PR.Panta Control and PR.Panta Analysis software (versions 1.6.3), respectively.

### 1.9 Cryo-EM sample preparation

The WT-ATP7B_Δ_ sample reconstituted in MSP1E3D1 for cryo-EM analysis was prepared as described above with the following modifications. Both 1 mM AlF_x_, prepared by pre-incubating AlCl_3_ with NaF in a 1:10 molar ratio for 10 min at RT, and 0.5 mM ADP were included in all sample preparation steps from the first anti-GFP resin wash. Monomeric ATP7B sample in DDM/CHS was directly after elution used for the reconstitution. Eluted ATP7B-MSP1E3D1 monomer was concentrated to ∼0.8 mg/mL (A_280_^0.1%^=1) and used for the grid preparation.

UltrAuFoil R1.2 / 1.3, 300 mesh gold grids were cleaned using GloQube Plus Glow discharge system (Quorum Technologies) with air gas, negative mode, 45 s glow, and 15 mA. Samples (3 µL per grid) were vitrified by plunge-freezing in liquid ethane using Vitrobot Mark IV System (Thermo Fisher Scientific) with wait time 10 s, blot force 0, and blot time 4.0 s, operated at 100% humidity and 4 °C.

### 1.10 Cryo-EM data acquisition

Cryo-EM data were collected at the UK national electron Bio-Imaging Centre (eBIC) on a Titan Krios 300 kV TEM (Thermo Fisher Scientific) equipped with a Bioquantum energy filter (Gatan), K3 direct electron detector (Gatan) in counted super-resolution mode, 50 µm C2 aperture and 100 µm objective aperture. The microscope was operated using EPU software version 3.6 (Thermo Fisher Scientific). The data was acquired at 130,000× magnification in aberration-free image shift (AFIS) mode with 0.645 Å calibrated pixel size. The set defocus ranged from 2.0 to 0.8 μm. Exposures of 50 e^-^/Å^2^ total dose were 1.4 s long with 50 frames, resulting in 1.0 e^-^/Å^2^ dose-per-frame.

### 1.11 Cryo-EM data processing

Data were processed using cryoSPARC^77^, version 4.6 and 4.7.0 (Structura Biotechnology). All jobs described below used the default settings except for the parameters listed. In total, 3,876 movies were imported and corrected for anisotropic full-frame motion with dose weighting^78^, followed by a patch-based contrast transfer function (CTF) estimation. After inspection, 2,913 exposures (75%) were accepted for blob picking with a minimum and maximum particle (ptc) diameters of 80 and 230 Å, respectively (1,511,231 ptcs; 519 picks per micrograph). After inspection, (1,100,550 ptcs; 378 P/mic.), particles were extracted with a 256-pixel sized box, Fourier-cropped to 128-pixel size (963,341 ptcs; 331 P/mic.). One round of 2D classification (200 classes, 2 final full iterations, 40 online-EM iterations, 400 batch size per class) followed by 2D class selection (33 classes selected; 334,845 ptcs ≈ 35%) provided particles for 3-class *ab initio* reconstruction. Subsequent heterogeneous refinement (Hetero-Ref) of the three generated volumes yielded one volume representing ATP7B_Δ_ in a nanodisc (4.42 Å; 204,593 ptcs). After non-uniform refinement^79^ (NU-Ref; minimize over per-particle scale True, reached 3.58 Å), this volume was used to generate 100 2D templates for picking (diameter 160 Å, 1,559,796 ptcs, 535 P/mic.). Inspected picks (1,165,777 ptcs, 400 P/mic.) were extracted (352 px → 128 px, 988,320 ptcs, 342 P/mic.) and used directly for 3D reconstruction by 3-class *ab initio* reconstruction followed by Hetero-Ref (6,000 batch size per class). Only one volume represented ATP7B_Δ_ in a nanodisc (4.84 Å, 508,873 ptcs). 2D classification of this volume (parameters as before + minimize over per-particle scale factors after first iteration True, force maximize over poses and shifts when aligning False) further refined the particle stack (160/200 selected; 467,238 ptcs ≈ 92%) before extraction (352 px; 466,812 ptcs; 2,877 mics; 162 P/mic.).

The particles were used for high-resolution heterogeneous *ab initio* reconstruction (HR-HAIR^80^) with three classes (3 Å maximum resolution, 6 Å initial resolution, Fourier radius step 0.005, centre structures in real space False, initial minibatch size 300, final minibatch size 1,000). Only one volume yielded ATP7B_Δ_ in a nanodisc, and this particle stack (261,233 ptcs) was used in NU-Ref reaching a resolution of 2.85 Å (enforce non-negativity True, batch size SNR factor 20, minimize over per-particle scale True, dynamic mask threshold 0.05, optimize per-group CTF parameters True with minimum fit resolution 7 Å). A mask covering the transmembrane domain and the A-, N-, and P-domains of ATP7B was generated and used for local refinement (Loc-Ref; use pose/shift gaussian prior during alignment True, enforce non-negativity True) and following 3D classification (4 classes, 3 Å filter resolution). After inspecting the volumes, focusing on difficult-to-resolve features (e.g. loops, N-domain, C-terminal tail), particles from three classes were combined and subjected to NU-Ref and Loc-Ref jobs as specified before, yielding a reconstruction at 2.79 Å (220,269 ptcs). Further sorting with a “subset particles by per-particle scale factor” job using three Gaussian distributions yielded a 150,030-ptc stack from the two higher-value subsets. Subsequent NU-Ref (parameters as before but optimize per-group CTF parameters True with minimum fit resolution 5 Å) and Loc-Ref (as before) resulted in a reconstruction at 2.80 Å with certain better-defined features, particularly the C-terminal tail. Local resolution estimation indicated a median resolution of 2.84 Å, with the central 50% of values ranging from 2.49 to 3.70 Å and the best-resolved regions reaching 1.86 Å. In the processing pipeline figure (Suppl. Fig. 3), we refer to this reconstruction as ATP7B_Δ_.

Throughout cryo-EM processing, weak density was consistently observed adjacent to the A-domain, suggesting partial occupancy by MBD6. Because this region did not resolve at high resolution, initial processing focused primarily on refinement of the ATP7B core. Only after introducing a 20Å-filter 3D classification to enrich particles containing MBD-like densities, we could iteratively refine both MBDs present in our construct in two configurations, which we designated the MBD5-up and MBD5-down states, respectively.

To obtain the MBD5-up state, the ATP7B_Δ_ volume was used as a reference for selecting 2D classes (200 classes, 3 Å maximum resolution, initial class uncertainty factor 1, circular mask diameter 160 Å, 80 online-EM iterations, 20 final full iterations, 400 batch size per class) generated from the original template-picked particles. Of the 255,402 particles selected (∼26%), *ab initio* reconstruction (4 classes, 3 Å maximum resolution, 6 Å initial resolution, 0.005 Fourier radius step; center in real space False; 300 initial minibatch; 100 final minibatch) yielded three protein-like volumes and one junk class. Particles from the protein-like classes (224,308 ptcs ≈ 88%) were subjected to NU-Ref (as above) and then to 3D classification (5 classes, 20 Å filter resolution). The resulting volumes were passed to Hetero-Ref (6,000 batch size per class) using the same particle stack. Approximately 20% of the particles (44,911), indicating MBD5 closer to the M domain, were NU-refined (20 Å initial lowpass resolution, enforce non-negativity True, minimize over per-particle scale True, optimize per-group CTF parameters True with minimum fit resolution 10 Å) to obtain a map containing both MBDs. To better trace the linker-region densities, we ran ten cycles of *ab initio* reconstruction (4 and later 2 classes, 20 Å filter resolution) followed by Hetero-Ref (2,000 batch size per class). This improved linker visibility but reduced global resolution. During these iterations, only the volumes containing better resolved MBDs were passed to subsequent rounds. The final 15,608-ptc stack was NU-refined (enforce non-negativity True, minimize over per-particle scale True, optimize per-group CTF parameters True with minimum fit resolution 10 Å), followed by Loc-Ref (mask excluding the nanodisc density with a fulcrum point placed on the MBD6 density, use pose/shift gaussian prior during alignment True, re-center shift each iteration True, minimize over per-particle scale True), yielding the MBD5-up state map at 4.41 Å resolution.

The MBD5-down state map was obtained using a similar processing strategy, but with an additional decoy-classification step. This required first obtaining a state in which MBD5 was located between the P- and A-domains. The same particle stack as previously used for the HR-HAIR described above was subjected to 6-class *ab initio* reconstruction (20 Å filter resolution) and Hetero-Ref (6,000 batch size per class), resulting in one 110,592-ptc (∼24 %) class with clear MBD5 density. Further 3D classification (5 classes, 20 Å filter resolution) sorted 45,014 particles (∼41%) into two volumes with prominent MBD5 density. This particle stack was NU-refined (enforce non-negativity True, minimize over per-particle scale True) to obtain the decoy volume for a Hetero-Ref (6,000 batch size per class) decoy-classification step. In this job, reference volumes representing a junk class, the previously obtained MBD5-up state, the ATP7B_Δ_, and a preliminary MBD5-up volume from the preceding NU-Ref were used to sort all template-picked particles. Particles assigned to the preliminary MBD5-up volume (121,359 ptcs ≈ 32%) were further 3D-classified (5 classes, 20 Å filter resolution). Three volumes showed a stronger MBD5 density (73,845 ptcs ≈ 61%). To avoid the bias introduced by the supervised decoy classification and to obtain a clean particle stack with well-defined MBDs and linkers, we ran 25 rounds of *ab initio* reconstruction (2 classes, 20–25 Å maximum resolution) followed by Hetero-Ref (2,000–3,000 batch size per class). At each iteration, particles from a volume with well-defined MBD5 density were carried forward, resulting in a final 12,960-ptc stack (∼18% of particles accepted from the initial 3D classification). Subsequent NU-Ref (enforce non-negativity True, minimize over per-particle scale True, optimize per-group CTF parameters True with minimum fit resolution 10 Å) and Loc-Ref (mask excluding the nanodisc density with a fulcrum point placed on the MBD6 density, use pose/shift gaussian prior during alignment True, enforce non-negativity True, minimize over per-particle scale True, NU filter order 3) yielded the MBD5-down state map at 4.09 Å resolution.

### 1.12 Model building

The initial model of ATP7B_Δ_ in an E2 conformation was obtained using AlphaFold 3 (AF3)^81^ accessed via https://alphafoldserver.com. The model was docked into the EM map using UCSF ChimeraX v1.9^82^ and Namdinator^83^ accessed via https://namdinator.au.dk. Iterative cycles of using Phenix real-space refinement^84^, Isolde v 1.10.1^85^, and Moorhen v0.22^86^ (https://moorhen.org) were used to build and refine the model. Because of the worse EM map quality, secondary structure restraints guided by the AlphaFold model template (https://alphafold.ebi.ac.uk/entry/AF-P35670-4-F1) were applied to the N-domain (residues 1034–1197) during adjustments using Isolde. The model was subjected to final real-space refinement in Phenix using Ramachandran restraints and secondary-structure restraints derived from the AlphaFold model. Residue numbering in the model provided is according to the canonical sequence with UniProt ID P35670-1. The final model consists of residues 643–1116 and 1142–1406. Images were prepared using ChimeraX v1.10-12^82^. Cryo-EM data collection, refinement, and validation statistics data are shown in Suppl. Table 1.

The final ATP7B_Δ_ model contains 11 cholesterol molecules (CLR), modelled based on cryo-EM density and molecular dynamics (MD) data. Initial density-guided docking was performed using the Cholesterol Docking ChimeraX plugin (github.com/TomasHeger/Cholesterol-Docking) with the protein-only model and cryo-EM map contoured at 0.141 (10.86σ). At the same contour level, the map was segmented using the Density Blob Classifier ChimeraX plugin (github.com/TomasHeger/Density-Blob-Classifier), with the protein-only model as a reference and a protein exclusion radius of 2.50 Å. Both workflows were implemented in Python using UCSF ChimeraX APIs^82^ . The automated cholesterol-docking workflow was developed independently, with its membrane-density detection strategy conceptually informed by published approaches for identifying membrane-associated density in cryo-EM maps, including MemBlob^87^ and SURFER^88^. Only CLRs docked into densities classified as “large” (68–120 Å³) were retained. Molecules occupying densities with obvious branching features, potentially indicative of other lipid species, were excluded. Initial refinement involved manual rotation of the molecules along their longitudinal axes to optimize their fit within the surrounding chemical environment, followed by refinement in Isolde. Further selection was guided by MD-derived occupancy data: only CLRs within 6 Å of residues with an average cholesterol occupancy above 20% across the four simulation runs were retained. This resulted in 11 CLRs, which were subsequently refined together with the rest of the model as described above.

Modelling of the low-resolution MBD5-up and MBD5-down configurations was based on the high-resolution ATP7B_Δ_ model. The corresponding cryo-EM maps were used to dock AF3-predicted models of MBD5 and MBD6. For the MBD5-down configuration, docking was guided by the previously published *Xt*ATP7B structure (PDB ID: 7SI3), whereas the solution NMR ensemble of the MBD5-6 construct served as a reference for the MBD5-up configuration ^30^. The connecting domain linkers were subsequently modelled using the Model Loops ChimeraX plugin^89^. Domain placement and linker geometry were refined in Isolde using AF3-derived secondary-structure restraints. During subsequent Phenix refinement, the regions shared with the high-resolution ATP7B_Δ_ model were restrained to preserve their secondary structure and sidechain rotamer conformations. The AF3-predicted models of MBD5 and MBD6 were also used as additional reference models during refinement. Detailed model validation statistics are summarized in Suppl. Table 1.

### 1.13 Structural bioinformatics

Multiple sequence alignment of human ATP7B orthologs (UniProt IDs: *Homo sapiens* P35670-1, *Pan troglodytes* H2Q7L5, *Mus musculus* Q64446, *Xenopus tropicalis* A0A6I8R0A5, *Danio rerio* A0A8N7T720, *Caenorhabditis elegans* G5EE14, *Oryza sativa* Q6H7M3, *Saccharomyces cerevisiae* P38995, *Legionella pneumophila* Q5ZWR1) was done using Jalview 2.11.5.1^90^ alignment tool with Clustal algorithm in default settings^91,92^.

The ATP7B sequence logo was generated via WebLogo 3 web application^93,94^ (www.weblogo.threeplusone.com) using an MSA input from ConservFold^95^ based on ColabFold v1.5.3^96^ using 95% similarity and E value >0.1 cutoff in the HHblits^97^ and MMseqs2 algorithms^98^. This MSA generation pipeline was run using human ATP7B (P35670-1) residues 484–1465. Residues were coloured by a colour-vision-deficiency accessible Clustal X colour scheme^99^ for MSA and hydrophobicity colouring for sequence logos^94^.

AlphaMissense^100^ pathogenicity score values for human ATP7B were downloaded from the AlphaFold Protein Structure Database (https://alphafold.ebi.ac.uk/, entry ID: AF-P35670-F1-v6)^101,102^. The associated heatmap was plotted using GraphPad Prism v10.3.1.

Cholesterol binding motif analysis was performed using the Cholesterol Binding Domain (CBD) Finder tool^44^ and human ATP7B (P35670-1) sequence as the input (www.mirna.in/tool/crac-carc-finder/). Only the CRAC (residues <u>709–720</u>, 949–959, 1073–1080, 1371–1378), CARC (residues 34–42, 527–536, 588–599, <u>648–659</u>, 758–770, <u>919–925</u>, <u>969–978</u>, 1238–1244, 1374–1381), and GXXXG (residues 43– 47, 280–284, 1099–1103, 1186–1190, 1281–1285) motifs localized in the M domain (<u>underlined</u>) were visualized. Lipid-like densities in the EM map are shown at a sigma level of 11.7. Molecular graphics and analyses were performed with UCSF ChimeraX v1.9-1.12^82^.

### 1.14 Molecular Dynamics Simulations

Coarse-grained molecular dynamics simulations were performed using the GROMACS simulation package (www.gromacs.org) (version 2025.2) together with the Martini 3 force field (martini3001 parameter set)^103^. The cryo-EM structure reported here was used as the wild-type protein model, from which the W939C mutant model was generated using the ChimeraX *swapaa* function with default settings. LipIDens^45^ was used for the protein preparation and simulation set-up. The protein structure was converted to a coarse-grained representation using martinize2 with automatic cysteine handling^104^. An elastic network model was applied to maintain the protein’s tertiary structure, using a force constant of 1000 kJ mol⁻¹ nm⁻² and lower and upper distance cutoffs of 0.5 nm and 0.9 nm, respectively^105^. The coarse-grained protein was embedded in a symmetric lipid bilayer generated using the LipIDens-modified version of insane^45,106^. The membrane composition consisted of POPC:DOPC:CHOL at a 40:40:20 molar ratio. The final system contained 270 POPC, 270 DOPC, and 135 cholesterol molecules. Parameters used were the standard Martini 3 parameter set^103^, except for cholesterol, where modified Martini 3 parameters were used^107^. The simulation box dimensions were 15 × 15 × 20 nm³. The system was solvated with 27,873 Martini 3 water beads and contained 406 Na⁺ and 406 Cl⁻ ions, corresponding to 0.15 M NaCl concentration in the simulation box. Energy minimization was first performed using the steepest-descent algorithm with 1000 steps. The system was subsequently equilibrated in two stages prior to the simulation. Four independent inputs with the same LipIDens settings were generated (Suppl. Fig. 12A) and four independent production run simulations were performed, each of 10 µs. Production run simulations were performed under the NPT ensemble at 310 K and 1 bar pressure, with a 20 fs timestep. Temperature coupling was performed with the v-rescale thermostat^108^ with protein, lipid and solvent coupled separately, and a temperature coupling constant of 1 ps. Pressure coupling was performed semi-isotropically, with the Parrinello-Rahman barostat^109^ and a pressure coupling time constant of 12 ps. A single cutoff of 1.1 nm was used for van der Waals interactions; for electrostatic interactions the reaction-field^110^ algorithm was used with a cutoff of 1.1 nm^111^. Lipid-protein interactions were analyzed using PyLipID^112^ by calculating lipid occupancy for each residue, defined as the frequency with which individual lipid species were observed in contact with a given protein residue over the course of the simulations.

## Supporting information

Supplementary File incl. supplementary Figures 1-13 and supplementary Tables 1-4.

## 2 Acknowledgements & Funding

We are grateful for excellent laboratory and administrative assistance by Anna Marie Nielsen and Karen Bech-Pedersen, for technical assistance by Taner Drace, Andreas Bøggild and Thomas Boesen of the Danish national cryo-EM facility EMBION, and by Jesper Karlsen and Rune Kidmose for resources and assistance with scientific computing and data storage at the EMCC data science facility at Aarhus University. We acknowledge Diamond for access and support of the cryo-EM facilities at the UK national electron Bio-Imaging Centre (eBIC), proposal BI37584. In this work, TH benefited from attending the Beginners Single Particle Cryo-EM course 2023 funded by the Wellcome Trust and MRC (218785/Z/19/Z). Molecular dynamics simulations were run at the Grendel cluster of the Centre for Scientific Computing Aarhus, supported by the Novo Nordisk Foundation equipment grant, ROBUST (NNF18OC0032608, NNF24OC0087976). Research in the ALD group is supported by grants from the Novo Nordisk Foundation (NNF23OC0079154) and the Independent Research Fund Denmark (DFF; 2098-00023B). NanoDSF measurements were performed at the Biophysics and Biochemistry Core Facility, Aarhus University, Denmark. This work was supported by an instrument grant from the Carlsberg Foundation (CF21-0164) to Magnus Kjærgaard. We thank Svetlana Lutsenko, Rasmus Kock Flygaard and Jessica Lu for valuable discussions. This work was supported by a professorship grant from the Lundbeck Foundation to PN (R310-2018-3713), an infrastructure grant from the Danish Ministry for Research and Higher Education (EMBION - 5072-00025B) and the Novo Nordisk Foundation (NNF20OC0060483), a MSCA postdoctoral fellowship grant to OM (101277193-WILSON-HORIZON-MSCA-2025-PF), an equipment grant from the Carlsberg Foundation (CF23-1394), and the CryoNet network supported by the Novo Nordisk Foundation.

## 3 Author contributions

TH – Conceptualization, Data curation, Formal analysis, Investigation, Methodology, Validation, Visualization, Writing original draft, Writing review and editing

OM – Data curation, Formal analysis, Investigation, Methodology, Validation, Visualization, Writing original draft, Writing review and editing

IMS – Formal analysis, Investigation

TD – Conceptualization, Methodology, Supervision, Writing review and editing

AD – Data curation, Formal analysis, Resources, Writing review and editing

PN – Funding acquisition, Resources, Supervision, Writing review and editing

CS – Conceptualization, Data curation, Formal analysis, Investigation, Methodology, Project administration, Supervision, Validation, Visualization, Writing original draft, Writing review and editing

## 4 Competing interests

The other authors declare no competing interests.

## 5 Data availability

The cryo-EM density maps have been deposited in the Electron Microscopy Data Bank (https://www.ebi.ac.uk/emdb) under the accession codes EMD-100822 (ATP7B_Δ_), EMD-100823 (MBD5-up) and EMD-100824 (MBD5-down). The atomic coordinates have been deposited in the Protein Data Bank (https://www.rcsb.org) under IDs 50PI (ATP7B_Δ_), 50PJ (MBD5-up) and 50PK (MBD5-down). CG-MD simulation files (.tpr, .gro, .xtc, .mdp), lipid occupancy and RMSD analysis results are accessible on Zenodo (WT-ATP7B simulation data: 10.5281/zenodo.21262460; ATP7B-W939C simulation data: 10.5281/zenodo.21526980). The Density Blob Classifier and Cholesterol Docking tools are available on GitHub (https://github.com/TomasHeger). Other data and materials used to support the findings of this study are available from the corresponding author upon reasonable request.

## Abbreviations

ATP7B: (copper-transporting ATPase 2)
CFTR: (cystic fibrosis transmembrane conductance regulator)
CG-MD: (coarse-grained molecular dynamics)
CHS: (cholesterol hemisuccinate)
CLR/CHOL: (cholesterol)
cryo-EM: (cryogenic electron microscopy)
MBD: (metal-binding domain)
NKA: (Na⁺/K⁺-ATPase)
*Os*: (*Oryza sativa*)
RMSD: (root mean square deviation)
SD: (standard deviation)
SERCA: (sarco/endoplasmic reticulum calcium ATPase)
WD: (Wilson disease)
*Xt*: (*Xenopus tropicalis*)

## Notes

### Competing Interest Statement

The authors have declared no competing interest.

## References

1. Scheiber, I. F., Mercer, J. F. B. & Dringen, R. Metabolism and functions of copper in brain. Prog. Neurobiol. 116, 33–57 (2014).

2. Chen, L., Min, J. & Wang, F. Copper homeostasis and cuproptosis in health and disease. Signal Transduct. Target. Ther. 7, 378 (2022).

3. Tümer, Z. & Møller, L. B. Menkes disease. Eur. J. Hum. Genet. 18, 511–518 (2010).

4. Członkowska, A. et al. Wilson disease. Nat. Rev. Dis. Primers 4, 21 (2018).

5. Dang, J. et al. Kidney involvement in Wilson’s disease: a review of the literature. Clin. Kidney J. 17, sfae058 (2024).

6. Dzieżyc-Jaworska, K., Litwin, T. & Członkowska, A. Clinical manifestations of Wilson disease in organs other than the liver and brain. Ann. Transl. Med. 7, S62 (2019).

7. Landrum, M. J. et al. ClinVar: public archive of relationships among sequence variation and human phenotype. Nucleic Acids Res. 42, D980–985 (2014).

8. Cooper, D., Ball, E. V. & Krawczak, M. The human gene mutation database. Nucleic Acids Res. 26, 285–287 (1998).

9. Sherry, S. T. et al. dbSNP: the NCBI database of genetic variation. Nucleic Acids Res. 29, 308–311 (2001).

10. Sandahl, T. D. et al. The Prevalence of Wilson’s Disease: An Update. Hepatology 71, 722–732 (2020).

11. Poujois, A. & Woimant, F. Challenges in the diagnosis of Wilson disease. Ann. Transl. Med. 7, S67 (2019).

12. Mohr, I. & Weiss, K. H. Current anti-copper therapies in management of Wilson disease. Ann. Transl. Med. 7, S69 (2019).

13. Overeem, A. W. et al. Pluripotent stem cell-derived bile canaliculi-forming hepatocytes to study genetic liver diseases involving hepatocyte polarity. J. Hepatol. 71, 344–356 (2019).

14. Concilli, M. et al. Pharmacoproteomics pinpoints HSP70 interaction for correction of the most frequent Wilson disease-causing mutant of ATP7B. Proc. Natl. Acad. Sci. U. S. A. 117, 32453–32463 (2020).

15. Chesi, G. et al. Identification of p38 MAPK and JNK as new targets for correction of Wilson disease-causing ATP7B mutants. Hepatology 63, 1842–1859 (2016).

16. Concilli, M., Iacobacci, S., Chesi, G., Carissimo, A. & Polishchuk, R. A systems biology approach reveals new endoplasmic reticulum-associated targets for the correction of the ATP7B mutant causing Wilson disease. Metallomics 8, 920–930 (2016).

17. Kerem, E. Mutation specific therapy in CF. Paediatr. Respir. Rev. 7 **Suppl 1**, S166–169 (2006).

18. Rowe, S. M. & Verkman, A. S. Cystic fibrosis transmembrane regulator correctors and potentiators. Cold Spring Harb. Perspect. Med. 3, a009761 (2013).

19. Palmgren, M. P-type ATPases: Many more enigmas left to solve. J. Biol. Chem. 299, 105352 (2023).

20. Stock, C. et al. Fast-forward on P-type ATPases: recent advances on structure and function. Biochem. Soc. Trans. 51, 1347–1360 (2023).

21. Wu, C.-C., Rice, W. J. & Stokes, D. L. Structure of a copper pump suggests a regulatory role for its metal-binding domain. Structure 16, 976–985 (2008).

22. Andersson, M. et al. Copper-transporting P-type ATPases use a unique ion-release pathway. Nat. Struct. Mol. Biol. 21, 43–48 (2014).

23. Gourdon, P. et al. Crystal structure of a copper-transporting PIB-type ATPase. Nature 475, 59–64 (2011).

24. Bitter, R. M. et al. Structure of the Wilson disease copper transporter ATP7B. Sci. Adv. 8, eabl5508 (2022).

25. Yang, G.-M. et al. Structures of the human Wilson disease copper transporter ATP7B. Cell Rep. 42, 112417 (2023).

26. Guo, Z. et al. Diverse roles of the metal binding domains and transport mechanism of copper transporting P-type ATPases. Nat. Commun. 15, 2690 (2024).

27. Ariöz, C., Li, Y. & Wittung-Stafshede, P. The six metal binding domains in human copper transporter, ATP7B: molecular biophysics and disease-causing mutations. Biometals 30, 823–840 (2017).

28. Huster, D. & Lutsenko, S. The distinct roles of the N-terminal copper-binding sites in regulation of catalytic activity of the Wilson’s disease protein. J. Biol. Chem. 278, 32212–32218 (2003).

29. Lutsenko, S., LeShane, E. S. & Shinde, U. Biochemical basis of regulation of human copper-transporting ATPases. Arch. Biochem. Biophys. 463, 134–148 (2007).

30. Achila, D. et al. Structure of human Wilson protein domains 5 and 6 and their interplay with domain 4 and the copper chaperone HAH1 in copper uptake. Proc. Natl. Acad. Sci. U. S. A. 103, 5729–5734 (2006).

31. Orädd, F., Steffen, J. H., Gourdon, P. & Andersson, M. Copper binding leads to increased dynamics in the regulatory N-terminal domain of full-length human copper transporter ATP7B. PLoS Comput. Biol. 18, e1010074 (2022).

32. Paweletz, L. C., et al. Anionic Phospholipids Stimulate the Proton Pumping Activity of the Plant Plasma Membrane P-Type H+-ATPase. Int. J. Mol. Sci. 24, 13106 (2023).

33. Brini, M. et al. Deletions and Mutations in the Acidic Lipid-binding Region of the Plasma Membrane Ca2+ Pump. Journal of Biological Chemistry 285, 30779–30791 (2010).

34. Cornelius, F., Habeck, M., Kanai, R., Toyoshima, C. & Karlish, S. J. D. General and specific lipid-protein interactions in Na,K-ATPase. Biochim. Biophys. Acta 1848, 1729–1743 (2015).

35. Dieudonné, T. et al. Autoinhibition and regulation by phosphoinositides of ATP8B1, a human lipid flippase associated with intrahepatic cholestatic disorders. Elife 11, e75272 (2022).

36. Purohit, R. et al. Cu+-specific CopB transporter: Revising P1B-type ATPase classification. Proc. Natl. Acad. Sci. U. S. A. 115, 2108–2113 (2018).

37. Autzen, H. E. et al. Interactions of a Bacterial Cu(I)-ATPase with a Complex Lipid Environment. Biochemistry 57, 4063–4073 (2018).

38. Braiterman, L. et al. Apical targeting and Golgi retention signals reside within a 9-amino acid sequence in the copper-ATPase, ATP7B. Am. J. Physiol. Gastrointest. Liver Physiol. 296, G433-444 (2009).

39. Guo, Y., Nyasae, L., Braiterman, L. T. & Hubbard, A. L. NH2-terminal signals in ATP7B Cu-ATPase mediate its Cu-dependent anterograde traffic in polarized hepatic cells. Am. J. Physiol. Gastrointest. Liver Physiol. 289, G904–916 (2005).

40. Leng, Y. et al. Long-Term Correction of Copper Metabolism in Wilson’s Disease Mice with AAV8 Vector Delivering Truncated ATP7B. Hum. Gene Ther. 30, 1494–1504 (2019).

41. Guerra, J. V. da S., et al. pyKVFinder: an efficient and integrable Python package for biomolecular cavity detection and characterization in data science. BMC Bioinformatics 22, 607 (2021).

42. Huster, D. et al. Diverse functional properties of Wilson disease ATP7B variants. Gastroenterology 142, 947–956.e5 (2012).

43. Schushan, M., Bhattacharjee, A., Ben-Tal, N. & Lutsenko, S. A structural model of the copper ATPase ATP7B to facilitate analysis of Wilson disease-causing mutations and studies of the transport mechanism. Metallomics 4, 669–678 (2012).

44. Sahu, V. K., Lokhande, K. B., Swamy, V. K., Basu, S. & Ranjan, A. Role of the transmembrane polar residues on CD151 in cholesterol binding. J. Biomol. Struct. Dyn. 43, 10268–10280 (2025).

45. Ansell, T. B. et al. LipIDens: simulation assisted interpretation of lipid densities in cryo-EM structures of membrane proteins. Nat. Commun. 14, 7774 (2023).

46. Pilankatta, R., Lewis, D., Adams, C. M. & Inesi, G. High yield heterologous expression of wild-type and mutant Cu+-ATPase (ATP7B, Wilson disease protein) for functional characterization of catalytic activity and serine residues undergoing copper-dependent phosphorylation. J. Biol. Chem. 284, 21307–21316 (2009).

47. Jensen, A.-M. L., Sørensen, T. L.-M., Olesen, C., Møller, J. V. & Nissen, P. Modulatory and catalytic modes of ATP binding by the calcium pump. EMBO J. 25, 2305–2314 (2006).

48. Clausen, J. D., Holdensen, A. N. & Andersen, J. P. Critical roles of interdomain interactions for modulatory ATP binding to sarcoplasmic reticulum Ca2+-ATPase. J. Biol. Chem. 289, 29123–29134 (2014).

49. Dmitriev, O. Y., Bhattacharjee, A., Nokhrin, S., Uhlemann, E.-M. E. & Lutsenko, S. Difference in stability of the N-domain underlies distinct intracellular properties of the E1064A and H1069Q mutants of copper-transporting ATPase ATP7B. J. Biol. Chem. 286, 16355–16362 (2011).

50. Morgan, C. T., Tsivkovskii, R., Kosinsky, Y. A., Efremov, R. G. & Lutsenko, S. The Distinct Functional Properties of the Nucleotide-binding Domain of ATP7B, the Human Copper-transporting ATPase. Journal of Biological Chemistry 279, 36363–36371 (2004).

51. Dmitriev, O. et al. Solution structure of the N-domain of Wilson disease protein: Distinct nucleotide-binding environment and effects of disease mutations. Proceedings of the National Academy of Sciences 103, 5302–5307 (2006).

52. Tsivkovskii, R., Efremov, R. G. & Lutsenko, S. The role of the invariant His-1069 in folding and function of the Wilson’s disease protein, the human copper-transporting ATPase ATP7B. J. Biol. Chem. 278, 13302–13308 (2003).

53. DiDonato, M., Hsu, H. F., Narindrasorasak, S., Que, L. & Sarkar, B. Copper-induced conformational changes in the N-terminal domain of the Wilson disease copper-transporting ATPase. Biochemistry 39, 1890–1896 (2000).

54. Chakraborty, K. et al. Wilson disease-causing mutations in the carboxyl terminus of ATP7B regulates its localization and Golgi exit selectively in the unpolarized cells. Metallomics 15, mfad051 (2023).

55. Braiterman, L. T. et al. Distinct phenotype of a Wilson disease mutation reveals a novel trafficking determinant in the copper transporter ATP7B. Proc. Natl. Acad. Sci. U. S. A. 111, E1364–1373 (2014).

56. Collins, C. J. et al. Direct Measurement of ATP7B Peptides Is Highly Effective in the Diagnosis of Wilson Disease. Gastroenterology 160, 2367–2382.e1 (2021).

57. Huster, D. et al. Defective cellular localization of mutant ATP7B in Wilson’s disease patients and hepatoma cell lines. Gastroenterology 124, 335–345 (2003).

58. Ferenci, P. et al. Age and Sex but Not ATP7B Genotype Effectively Influence the Clinical Phenotype of Wilson Disease. Hepatology 69, 1464–1476 (2019).

59. Dode, L. et al. Functional Comparison between Secretory Pathway Ca2+/Mn2+-ATPase (SPCA) 1 and Sarcoplasmic Reticulum Ca2+-ATPase (SERCA) 1 Isoforms by Steady-state and Transient Kinetic Analyses. Journal of Biological Chemistry 280, 39124–39134 (2005).

60. Christensen, M. E. et al. Active conformations of neuronal Na+, K+-ATPase isoforms and a disease-causing mutant. Nat. Commun. 17, 9278 (2026).

61. Hordyjewska, A., Popiołek, Ł. & Kocot, J. The many ‘faces’ of copper in medicine and treatment. Biometals 27, 611–621 (2014).

62. Linder, M. C. & Hazegh-Azam, M. Copper biochemistry and molecular biology. Am. J. Clin. Nutr. 63, 797S–811S (1996).

63. Calvo, J. S. et al. Functional Screen of Wilson Disease ATP7B Variants Reveals Residual Transport Activities. Hum. Mutat. 2025, 7485658 (2025).

64. Forbes, J. R. & Cox, D. W. Copper-dependent trafficking of Wilson disease mutant ATP7B proteins. Hum. Mol. Genet. 9, 1927–1935 (2000).

65. Forbes, J. R. & Cox, D. W. Functional Characterization of Missense Mutations in ATP7B: Wilson Disease Mutation or Normal Variant? The American Journal of Human Genetics 63, 1663–1674 (1998).

66. Zhu, M., Dong, Y., Ni, W. & Wu, Z.-Y. Defective roles of ATP7B missense mutations in cellular copper tolerance and copper excretion. Mol. Cell. Neurosci. 67, 31–36 (2015).

67. van den Berghe, P. V. E., et al. Reduced expression of ATP7B affected by Wilson disease-causing mutations is rescued by pharmacological folding chaperones 4-phenylbutyrate and curcumin. Hepatology 50, 1783–1795 (2009).

68. Fiedorczuk, K. & Chen, J. Mechanism of CFTR correction by type I folding correctors. Cell 185, 158–168.e11 (2022).

69. Payne, A. S., Kelly, E. J. & Gitlin, J. D. Functional expression of the Wilson disease protein reveals mislocalization and impaired copper-dependent trafficking of the common H1069Q mutation. Proc. Natl. Acad. Sci. U. S. A. 95, 10854–10859 (1998).

70. Parisi, S. et al. Characterization of the most frequent ATP7B mutation causing Wilson disease in hepatocytes from patient induced pluripotent stem cells. Sci. Rep. 8, 6247 (2018).

71. Denisov, I. G., Baas, B. J., Grinkova, Y. V. & Sligar, S. G. Cooperativity in cytochrome P450 3A4: linkages in substrate binding, spin state, uncoupling, and product formation. J. Biol. Chem. 282, 7066–7076 (2007).

72. Sumida, K. H. et al. Improving Protein Expression, Stability, and Function with ProteinMPNN. J. Am. Chem. Soc. 146, 2054–2061 (2024).

73. Gietz, R. D., Schiestl, R. H., Willems, A. R. & Woods, R. A. Studies on the transformation of intact yeast cells by the LiAc/SS-DNA/PEG procedure. Yeast 11, 355–360 (1995).

74. Jidenko, M., Lenoir, G., Fuentes, J. M., le Maire, M. & Jaxel, C. Expression in yeast and purification of a membrane protein, SERCA1a, using a biotinylated acceptor domain. Protein Expr. Purif. 48, 32–42 (2006).

75. Cong, A. T. Q., Witter, T. L. & Schellenberg, M. J. High-efficiency recombinant protein purification using mCherry and YFP nanobody affinity matrices. Protein Sci. 31, e4383 (2022).

76. Heger, T. et al. eGFP as an All-in-One Tag for Purification of Membrane Proteins. Methods in Molecular Biology 2652, 171–186 (2023).

77. Punjani, A., Rubinstein, J. L., Fleet, D. J. & Brubaker, M. A. cryoSPARC: algorithms for rapid unsupervised cryo-EM structure determination. Nat. Methods 14, 290–296 (2017).

78. Grant, T. & Grigorieff, N. Measuring the optimal exposure for single particle cryo-EM using a 2.6 Å reconstruction of rotavirus VP6. Elife 4, e06980 (2015).

79. Punjani, A., Zhang, H. & Fleet, D. J. Non-uniform refinement: adaptive regularization improves single-particle cryo-EM reconstruction. Nat. Methods 17, 1214–1221 (2020).

80. Kim, K., Li, H. & Clarke, O. B. High-resolution ab initio reconstruction enables cryo-EM structure determination of small particles. *bioRxiv* Preprint at 10.1101/2025.09.08.674935 (2025).

81. Abramson, J. et al. Accurate structure prediction of biomolecular interactions with AlphaFold 3. Nature 630, 493–500 (2024).

82. Meng, E. C. et al. UCSF ChimeraX: Tools for structure building and analysis. Protein Sci. 32, e4792 (2023).

83. Kidmose, R. T. et al. Namdinator - automatic molecular dynamics flexible fitting of structural models into cryo-EM and crystallography experimental maps. IUCrJ 6, 526–531 (2019).

84. Liebschner, D. et al. Macromolecular structure determination using X-rays, neutrons and electrons: recent developments in Phenix. Acta Crystallogr. D Struct. Biol. 75, 861–877 (2019).

85. Croll, T. I. ISOLDE: a physically realistic environment for model building into low-resolution electron-density maps. Acta Crystallogr. D Struct. Biol. 74, 519–530 (2018).

86. Sanchez, F. Moorhen: a web browser molecular graphics program based on the Coot desktop program. Preprint at https://moorhen.org/ (2025).

87. Csizmadia, G., Farkas, B., Katona, E., Tusnády, G. E. & Hegedűs, T. Using MemBlob to Analyze Transmembrane Regions Based on Cryo-EM Maps. in 123–130 (2020). doi:10.1007/978-1-0716-0270-6_9.

88. Bharadwaj, A., Veerbeek, L. & Jakobi, A. Interactive segmentation of membrane and membrane-mimic densities in cryo-EM maps. Acta Crystallogr. D Struct. Biol. 82, 848–861 (2026).

89. Eswar, N. et al. Comparative protein structure modeling using Modeller. Curr. Protoc. Bioinformatics **Chapter 5**, Unit-5.6 (2006).

90. Waterhouse, A. M., Procter, J. B., Martin, D. M. A., Clamp, M. & Barton, G. J. Jalview Version 2--a multiple sequence alignment editor and analysis workbench. Bioinformatics 25, 1189–1191 (2009).

91. Troshin, P. V, Procter, J. B. & Barton, G. J. Java bioinformatics analysis web services for multiple sequence alignment--JABAWS:MSA. Bioinformatics 27, 2001–2 (2011).

92. Troshin, P. V et al. JABAWS 2.2 distributed web services for Bioinformatics: protein disorder, conservation and RNA secondary structure. Bioinformatics 34, 1939–1940 (2018).

93. Schneider, T. D. & Stephens, R. M. Sequence logos: a new way to display consensus sequences. Nucleic Acids Res. 18, 6097–6100 (1990).

94. Crooks, G. E., Hon, G., Chandonia, J.-M. & Brenner, S. E. WebLogo: a sequence logo generator. Genome Res. 14, 1188–1190 (2004).

95. Graham, C., Stansfeld, P. & Rodriguesp, C. ConservFold: Conservation to 3D structure generator. Preprint at https://zenodo.org/records/10062829 (2023).

96. Mirdita, M. et al. ColabFold: making protein folding accessible to all. Nat. Methods 19, 679–682 (2022).

97. Remmert, M., Biegert, A., Hauser, A. & Söding, J. HHblits: lightning-fast iterative protein sequence searching by HMM-HMM alignment. Nat. Methods 9, 173–175 (2011).

98. Steinegger, M. & Söding, J. MMseqs2 enables sensitive protein sequence searching for the analysis of massive data sets. Nat. Biotechnol. 35, 1026–1028 (2017).

99. Larkin, M. A. et al. Clustal W and Clustal X version 2.0. Bioinformatics 23, 2947–2948 (2007).

100. Cheng, J. et al. Accurate proteome-wide missense variant effect prediction with AlphaMissense. Science 381, eadg7492 (2023).

101. Jumper, J. et al. Highly accurate protein structure prediction with AlphaFold. Nature 596, 583–589 (2021).

102. Bertoni, D. et al. AlphaFold Protein Structure Database 2025: a redesigned interface and updated structural coverage. Nucleic Acids Res. 54, D358–D362 (2026).

103. Souza, P. C. T. et al. Martini 3: a general purpose force field for coarse-grained molecular dynamics. Nat. Methods 18, 382–388 (2021).

104. Kroon, P. C. et al. Martinize2 and Vermouth provide a unified framework for molecular topology generation. Elife 12, RP90627 (2025).

105. Periole, X., Cavalli, M., Marrink, S.-J. & Ceruso, M. A. Combining an Elastic Network With a Coarse-Grained Molecular Force Field: Structure, Dynamics, and Intermolecular Recognition. J. Chem. Theory Comput. 5, 2531–2543 (2009).

106. Wassenaar, T. A., Ingólfsson, H. I., Böckmann, R. A., Tieleman, D. P. & Marrink, S. J. Computational Lipidomics with insane: A Versatile Tool for Generating Custom Membranes for Molecular Simulations. J. Chem. Theory Comput. 11, 2144–2155 (2015).

107. Borges-Araújo, L. et al. Martini 3 Coarse-Grained Force Field for Cholesterol. J. Chem. Theory Comput. 19, 7387–7404 (2023).

108. Bussi, G., Donadio, D. & Parrinello, M. Canonical sampling through velocity rescaling. J. Chem. Phys. 126, 014101 (2007).

109. Parrinello, M. & Rahman, A. Polymorphic transitions in single crystals: A new molecular dynamics method. J. Appl. Phys. 52, 7182–7190 (1981).

110. Barker, J. A. & Watts, R. O. Monte Carlo studies of the dielectric properties of water-like models. Mol. Phys. 26, 789–792 (1973).

111. de Jong, D. H., Baoukina, S., Ingólfsson, H. I. & Marrink, S. J. Martini straight: Boosting performance using a shorter cutoff and GPUs. Comput. Phys. Commun. 199, 1–7 (2016).

112. Song, W. et al. PyLipID: A Python Package for Analysis of Protein-Lipid Interactions from Molecular Dynamics Simulations. J. Chem. Theory Comput. 18, 1188–1201 (2022).

