## Supplementary File incl. supplementary Figures 1-13 and supplementary Tables 1-4. for "Wilson disease mechanisms manipulate ATP7B activity"

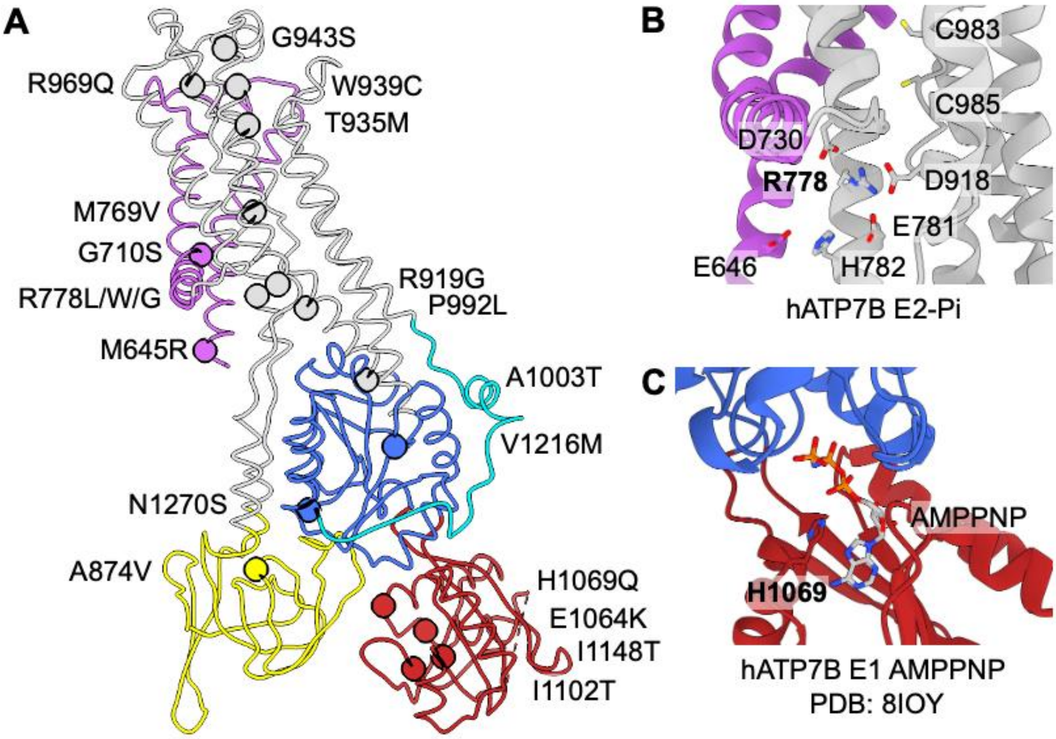


**Suppl. Fig. 1: Common WD mutations (single nucleotide polymorphisms, SNPs) and molecular mechanisms of WD mutations R778L and H1069Q.**

A) SNPs are localized throughout ATP7B. Depicted are the most prevalent variants as well as R969Q and W939C, the focus variants of this study.

B) Variations of R778 (L/W/G) are most common in Asia, and lead to decreased copper transport activity and retention of ATP7B in the ER [PMID: 18203200, 28842499, 9837819]. In our structure R778 sits in the centre of a 4-residue negatively charged ring at the ion entry site, occluding the cytoplasmic ion entry pathway.

C) The most common WD variant in Europe H1069Q localizes to the N-domain. Histidine coordinates the alpha-phosphate of bound nucleotides, and the mutant causes reduced phosphorylation, decreased copper transport and ATPase activity, and improper pump localization and trafficking [PMID: 9261163, 9311736, 9724794, 12557139, 19937698, 22240481, 23843626, 26660341, 29674751, 30965071]. Furthermore, a H1069Q substitution reduces nucleotide affinity and destabilizes the N-domain [PMID: 12551905,15205462,16567646,21398519].

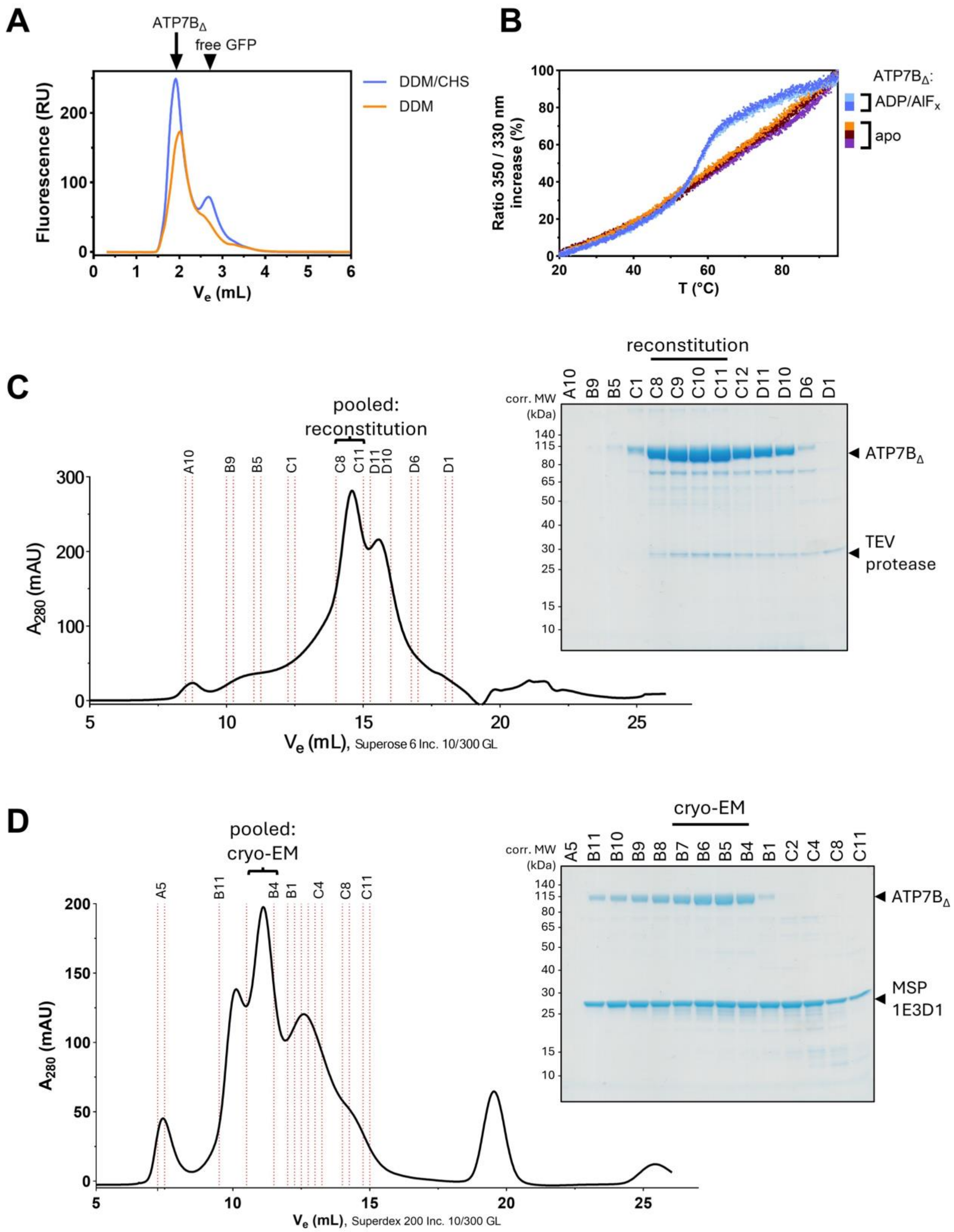


**Suppl. Fig. 2: $ATP7B_{\Delta}$ production and reconstitution for structural and functional studies.**

A) Fluorescence trace chromatogram ($\lambda_{ex}$ 488 nm / $\lambda_{em}$ 512 nm) from FSEC analysis of $ATP7B_{\Delta}$ crude yeast membranes (5 mg/ml total protein) solubilized in 1.25% (w/v) DDM without and with 0.25% CHS. A higher solubilization efficiency for DDM/CHS is observed. We previously published the results shown in subpanel A) as part of Figure 3 in Heger T. *et al.* (2023).

B) Nano differential scanning fluorimetry (nanoDSF) analysis of purified $ATP7B_{\Delta}$ in DDM/CHS. The addition of 0.5 mM ADP and 1 mM $AlF_x$ during purification stabilizes the sample, a clear transition in the melting curve is apparent.

C) Size exclusion chromatography and accompanying SDS-PAGE of purified $ATP7B_{\Delta}$ in DDM/CHS with 0.5 mM ADP and 1 mM $AlF_x$. Fractions pooled for the subsequent reconstitution are marked.

D) Size exclusion chromatography and accompanying SDS-PAGE of $ATP7B_{\Delta}$ in MSP1E3D1 nanodiscs with 0.5 mM ADP and 1 mM $AlF_x$. Fractions pooled for cryo-EM grid preparation are marked.

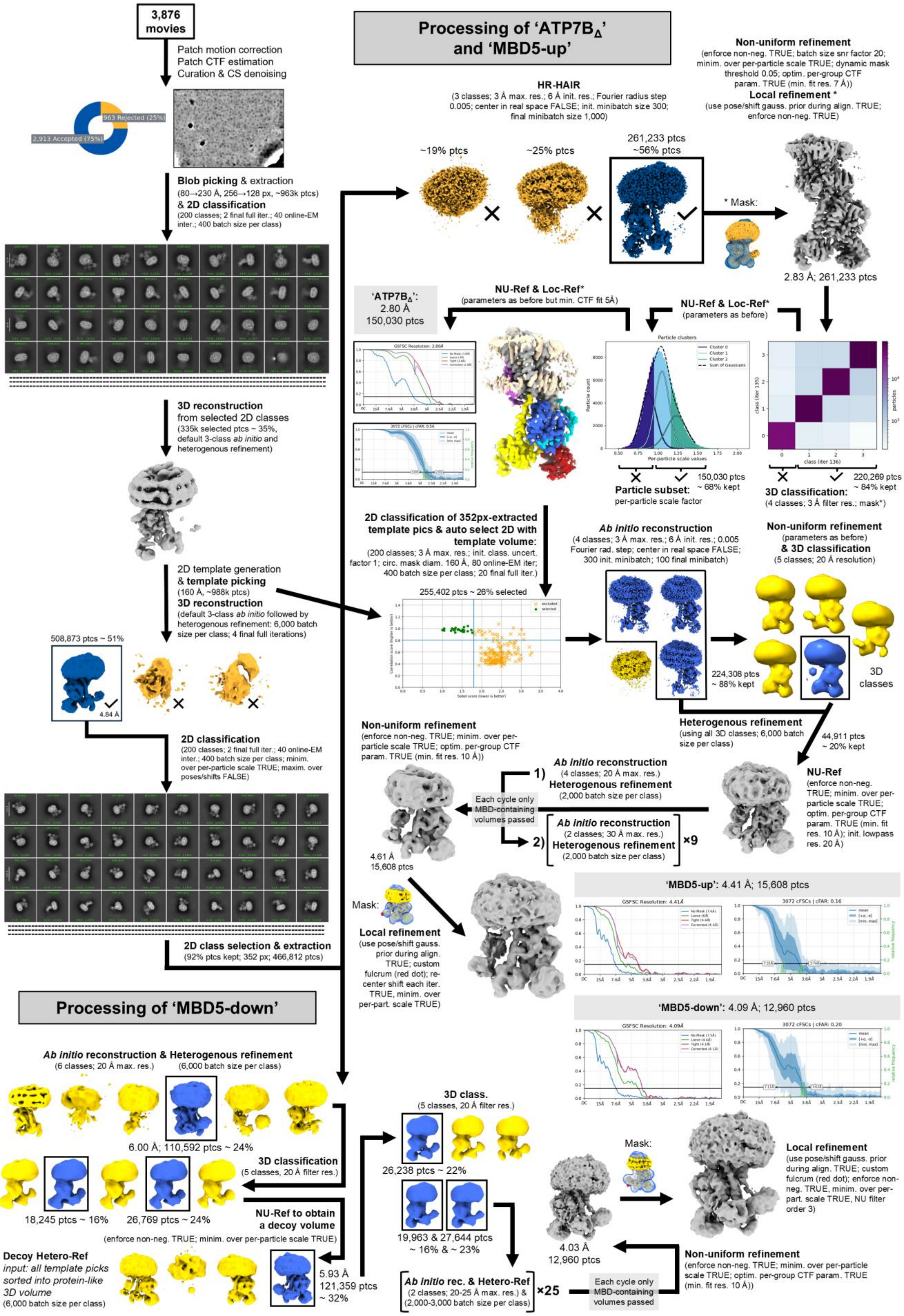


**Suppl. Fig. 3: Cryo-EM processing workflows, including a representative denoised micrograph, 2D classes, graphs for GS-FSC and cFAR scores, as well as the final cryo-EM maps.**

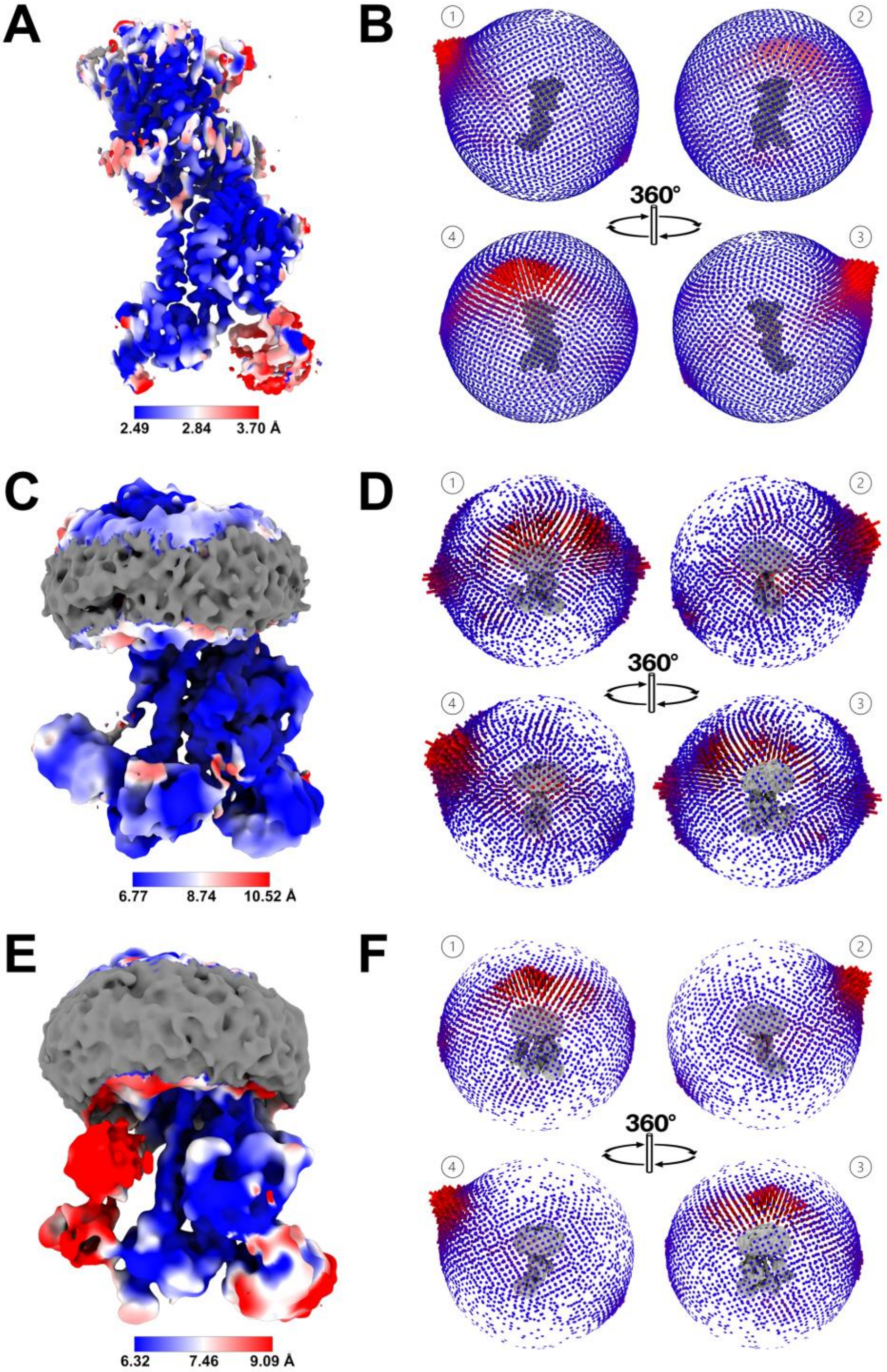


**Suppl. Fig. 4: Cryo-EM map assessment.**

A, C, E) Obtained cryo-EM maps coloured by local resolution scale (25$^{th}$-75$^{th}$ resolution percentile range). Maps represent the following structures: A - $ATP7B_{\Delta}$ (sigma 10.01), C - MBD5-down (sigma 5.65), and E - MBD5-up (sigma 4.91). Density outside the mask used for local resolution estimation is coloured in grey.

B, D, F) Three-dimensional particle orientation distribution plots for the corresponding maps on the left.

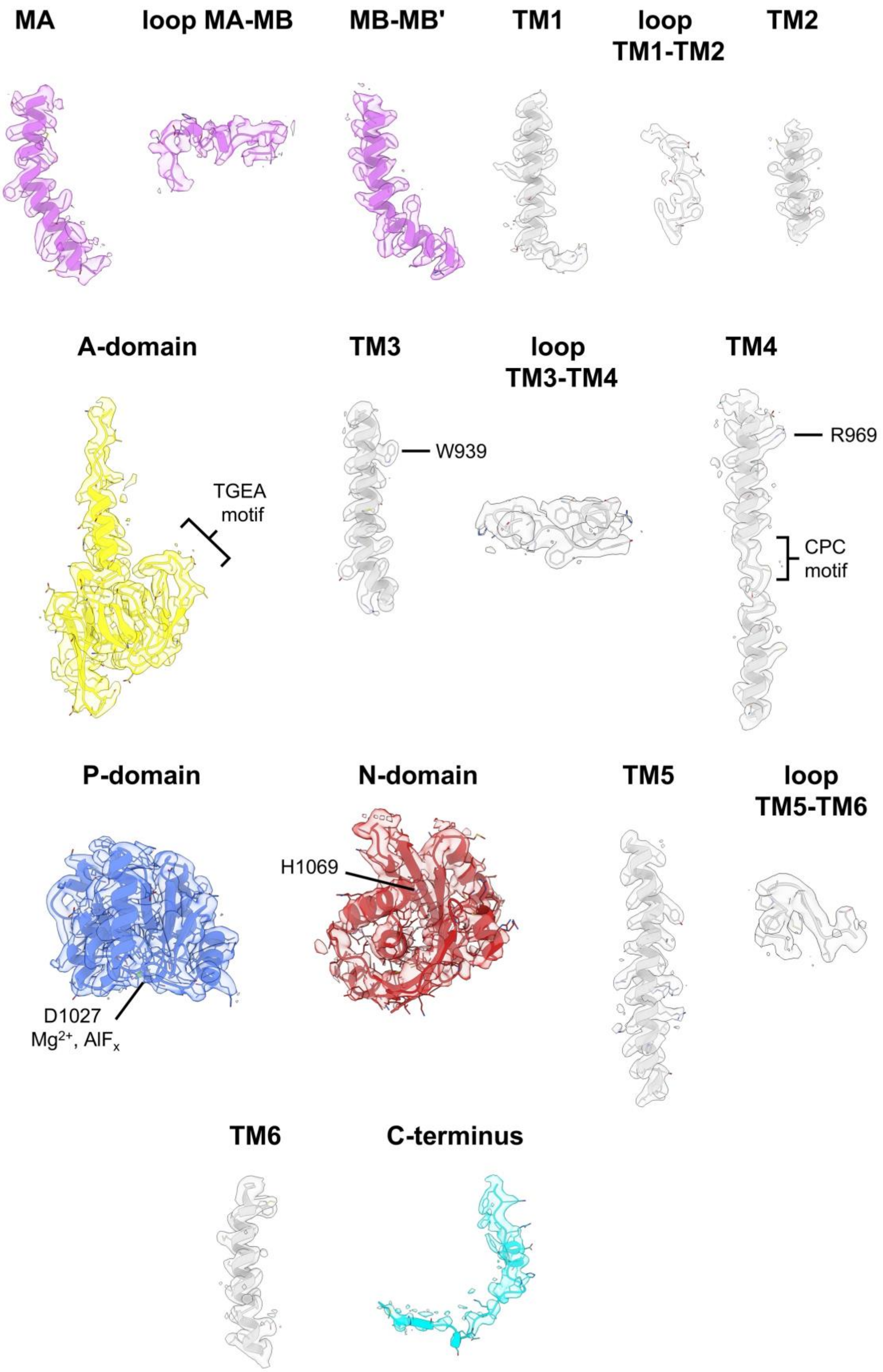


**Suppl. Fig. 5: Map to model fit ATP7B$_{\Delta}$: cryo-EM densities of ATP7B moieties in sequential order (sigma 7.7). Conserved motifs and mutations in one-letter code.**

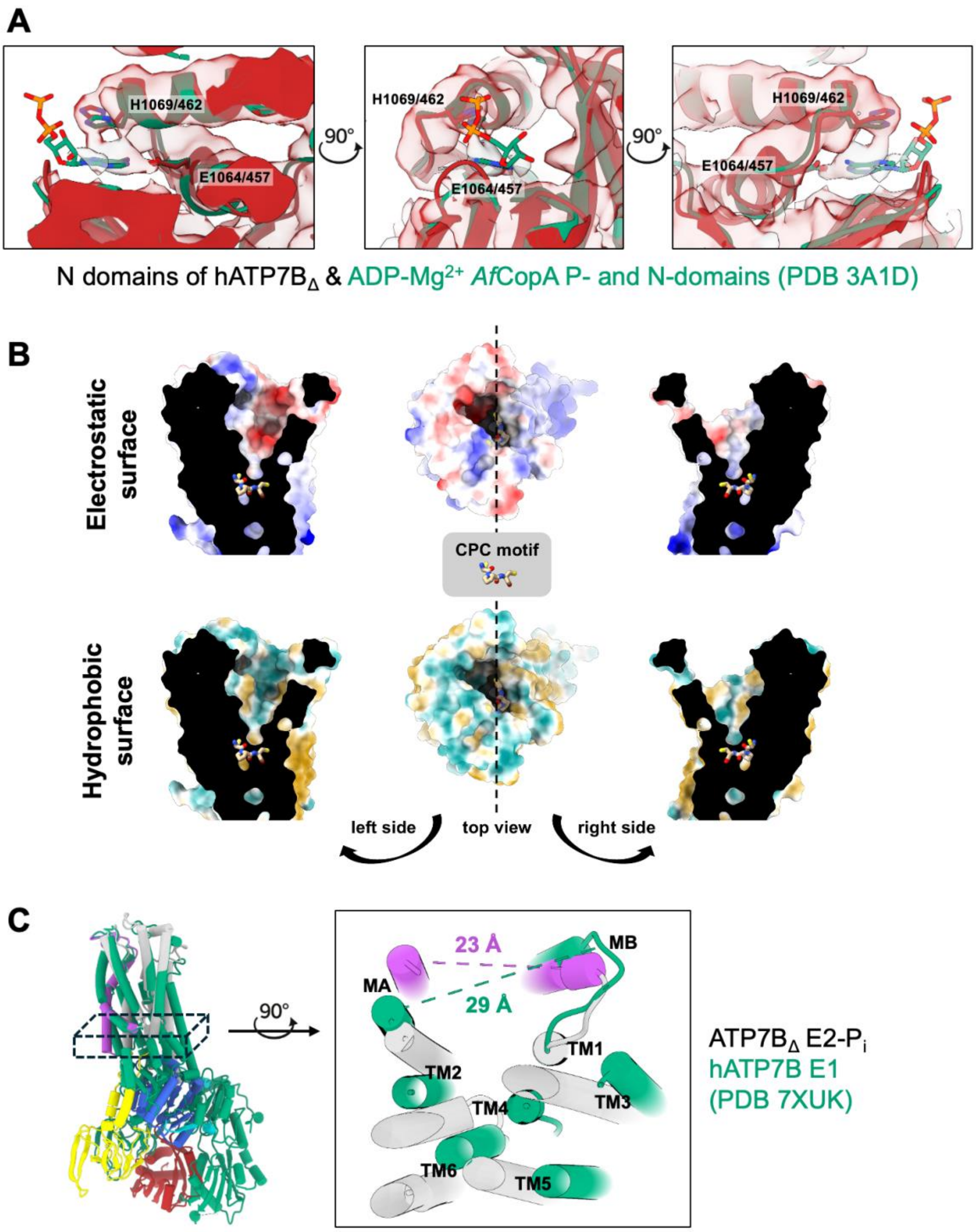


**Suppl. Fig. 6: Structural features of hATP7B$_{\Delta}$.**

A) Overlay of our human ATP7B$_{\Delta}$ map and model (no ADP modeled) and a crystal structure of *Af*CopA N- and P-domains (teal, PDB 3A1D, including ADP) in three 90º offset views. Clear partial overlap between our map and the ADP from the crystal structure can be observed. Conserved E1069/457 (ATP7B/*Af*CopA) coordinates the adenine ring, while WD residue H1069/462 interacts with the alpha phosphate of the nucleotide. Notably, H1069 occupies a spatially analogous position to SERCA E439, which contributes to modulatory ATP binding through a Mg$^{2+}$-mediated interaction.

B) Top: Electrostatic tunnel surface. The molecular surface was coloured according to Coulombic electrostatic potential using a red-white-blue colour scale (-10 to +10 kcal·mol$^{-1}$·e$^{-1}$), with blue representing positive electrostatic potential and red representing negative electrostatic potential. Bottom: Hydrophobic tunnel surface. The molecular surface was coloured according to the Fauchère molecular lipophilicity potential (MLP), with dark goldenrod representing positive (lipophilic) values and dark cyan representing negative (hydrophilic) values.

C) Overlay of our human ATP7B$_{\Delta}$ model in an E2-Pi conformation and hATP7B in an E1 conformation (PDB 7XUK) revealing a wider space between MB' and MA in the E1 state, allowing for docking of a copper delivery system.

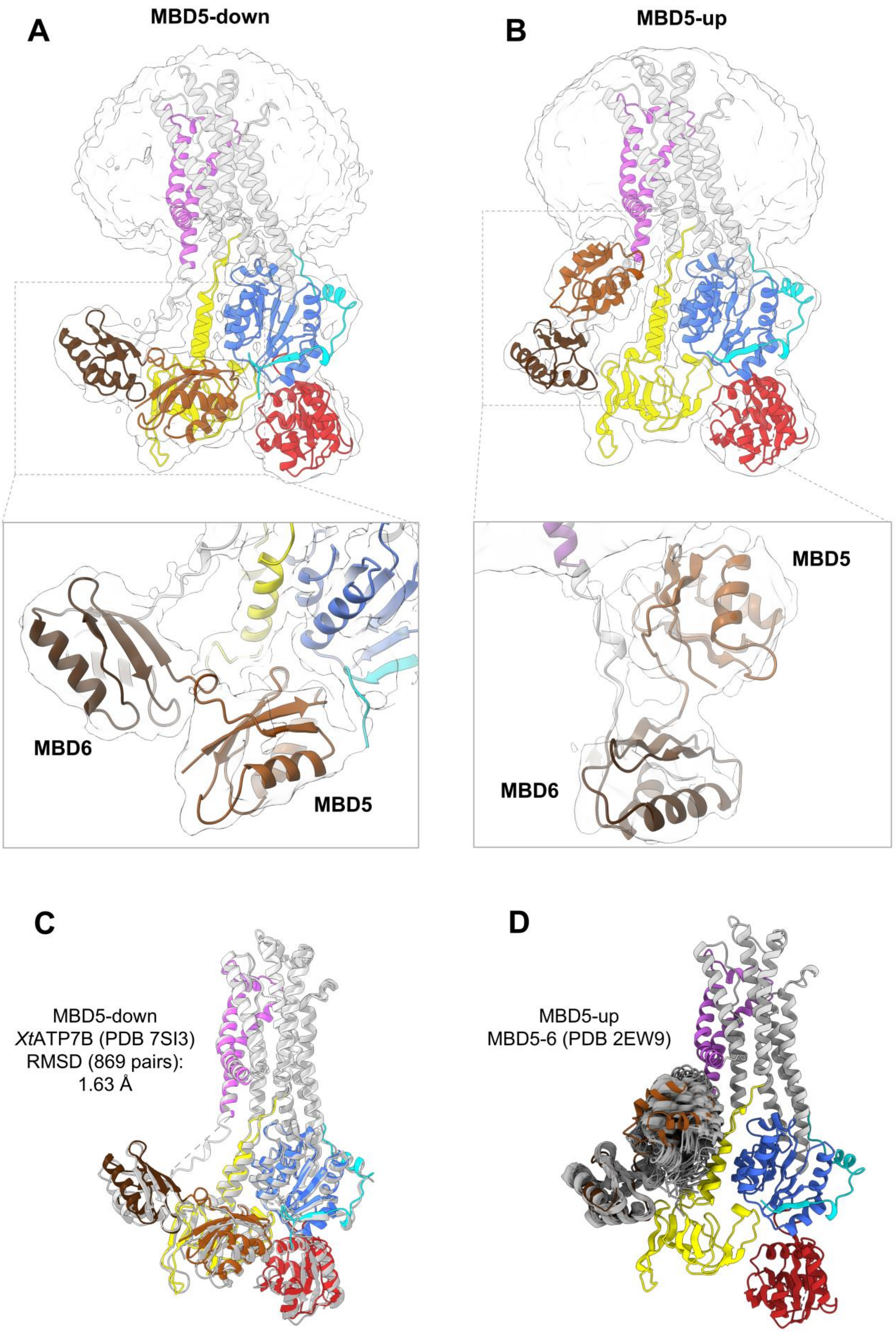


**Suppl. Fig. 7: Varied localization of MBD5/6 for priming of copper delivery to $ATP7B_{\Delta}$ in the E2-$P_i$ conformation.**

A) Map and docked model of the MBD5-down state (sigma level 5.3). While MBD6 is posed on the A-domain, MBD5 is docked between the A- and P-domains, where a hydrophobic loop of MBD5 (residues 523–526) faces a hydrophobic cavity formed between the structured A-domain linker and the P-domain. This interaction is furthered by the C-terminal tail (residues 1406-1410) contacting MBD5 adjacent to its $^{499}$CASC$^{502}$ copper-binding motif. These poses of MBD5/6 are similar to the ones described by Bitter *et al.*, for *Xt*ATP7B in an E2-$P_i$ conformation.
Below: close-up view of the map and model in the MBD5-down pose.

B) Map and docked model of the MBD5-up state (sigma level 5.0). MBD5 moves upwards towards MB' by about 52°, while MBD6 remains atop the A-domain. Below: close-up view of the map and model in the MBD5-up pose. MBD5 interacts with the MBD6-MA linker and is closer to the MB' linker than in the MBD5-down pose.

C) Structural comparison of our MBD5-down state and *Xt*ATP7B (PDB 7SI3), both in an E2-P*i* conformation.

D) Overlay of solution NMR structure ensemble of MBD5-MBD6 (PDB ID: 2EW9) with MBD5-up state, aligned on MBD6. While the relative orientation of the MBDs matches well with the solution NMR structure, the NMR ensemble suggests a degree of bending between the domains that would be consistent with the lower quality and broader MBD5 density in our map compared to MBD5 density in the MBD5-down pose.

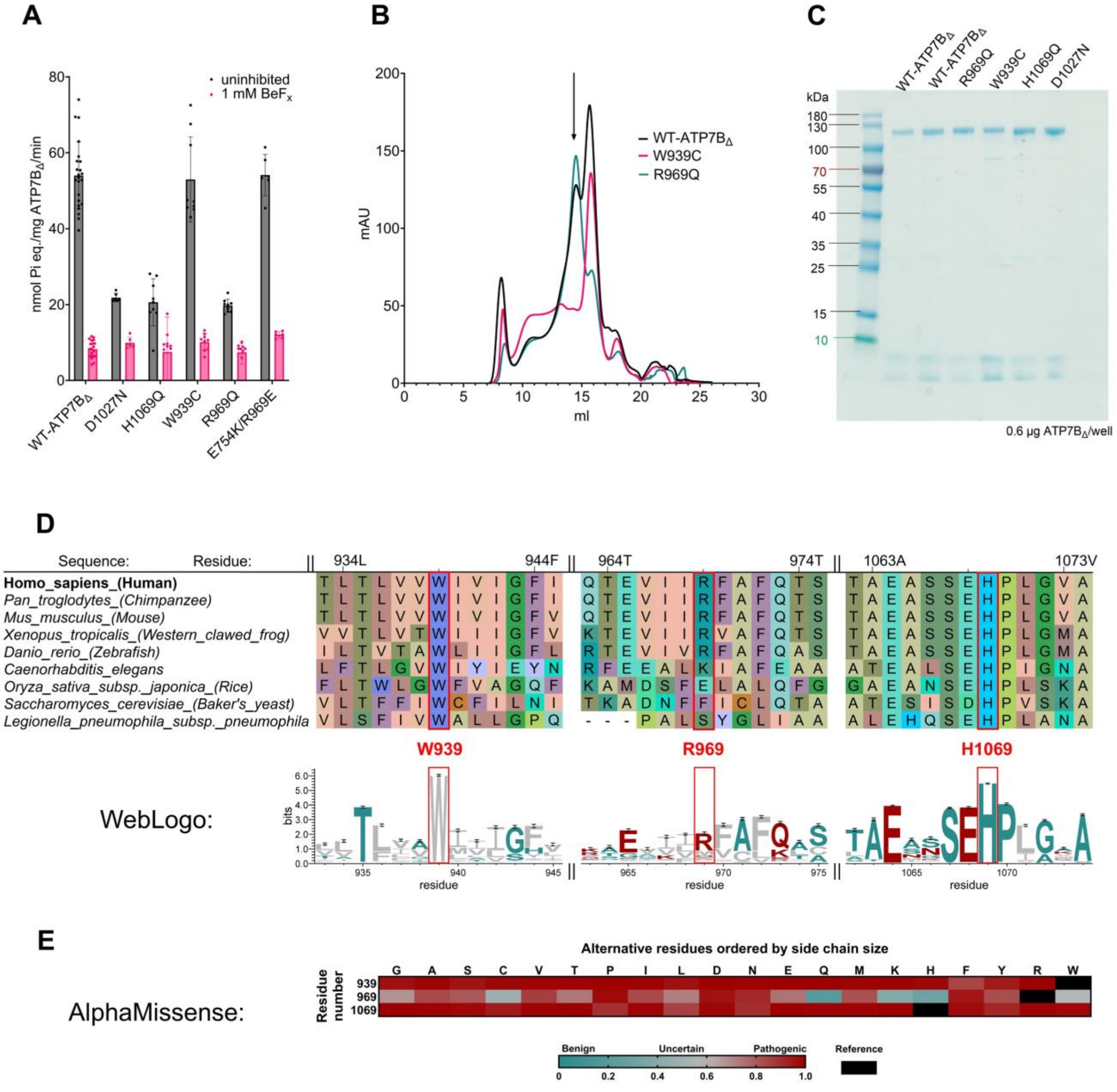


**Suppl. Fig. 8: Further analysis of ATP7B variants.**

A) ATPase activity assay of ATP7B$_\Delta$. Catalytically inactive (D1027N), three WD mutations and the double mutant E754K/R969E in the absence and presence of the inhibitor BeF$_x$. Absolute ATPase activity values are given in nmol $P_i$ / mg ATP7B$_\Delta$ / min.

B) SEC run comparison of DDM/CHS-solubilized and purified ATP7B$_\Delta$ and mutants W939C and R969Q. The arrow marks the monomeric fraction of ATP7B$_\Delta$ and its variants loaded onto the gel in C.

C) SDS-PAGE analysis of purified ATP7B$_\Delta$ samples.

D) Multiple sequence alignment of ATP7B homologues and WebLogo conservation plot. The views are condensed around the three WD mutant residues compared in this manuscript: W939, R969 and H1069. While W939 and 1069 are highly conserved, R969 seems to be maintained only in higher eukaryotes.

E) AlphaMissense plot for the three residues W939, R969 and H1069.

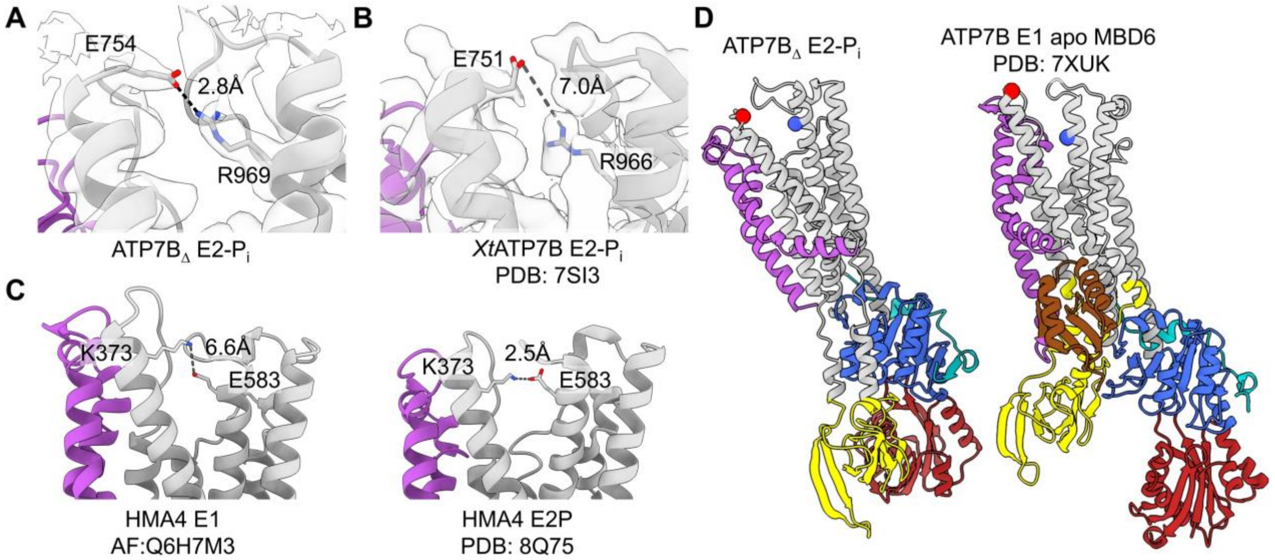


**Suppl. Fig 9: Conserved salt bridge stabilizes E2 conformation of ATP7B.**

A) Model and map (sigma 8.4) of human ATP7B around the salt bridge between E754 and R969 (distance OE-NH 2.8Å) in an E2-P$_i$ conformation.

B) Model and map (sigma 1.1) of *Xenopus tropicalis* XtATP7B in an E2-P$_i$ conformation around the salt bridge between E754 and R969 (distance OE-NH 7.0Å) due to different model building.

C) Models of HMA4 from *Oryza sativa* in E1 (AlphaFold DB: AF-Q6H7M3-F1-v6) and E2 (PDB:8Q75) conformations with charge-inversed salt bridge between K373 and E583. Distances between NZ and OE 2.5Å (E2P) vs. 6.6Å (E1).

D) MBD6 binding below MB' and A-domain rotation cause breaking of salt bridge in the E2 to E1 transition.

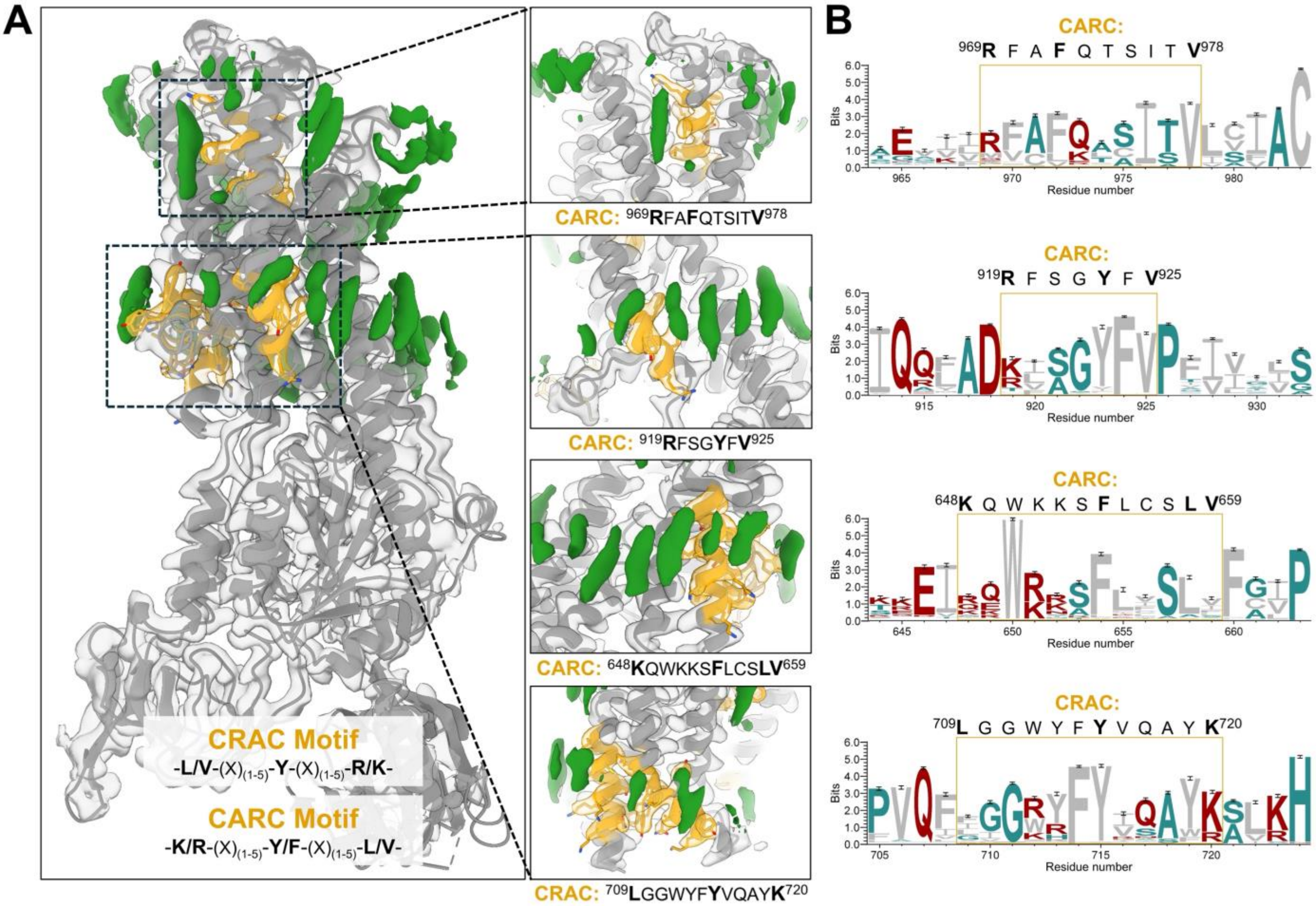


**Suppl. Fig 10: Cholesterol binding site analysis of the CRAC and CARC motifs found in hATP7B and putative lipid densities in the cryo-EM map.**

A) Visualization of lipid-like densities (green) in our cryo-EM map (Sigma 11.7), nearby annotated cholesterol-binding CRAC and CARC motifs are shown in stick representation (gold).

B) Conservation analysis of the identified CRAC and CARC motifs using WebLogo in standard colouring.

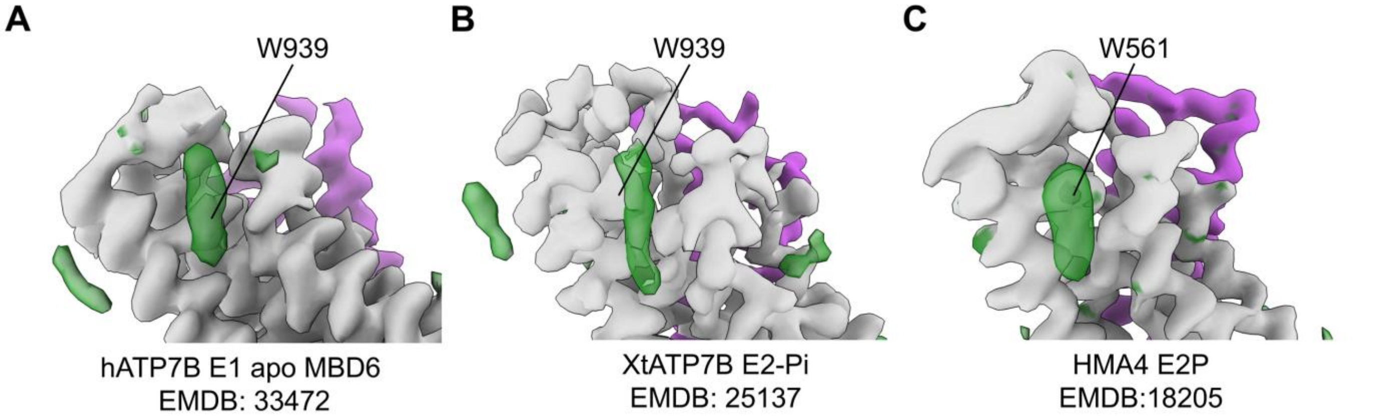


**Suppl. Fig 11: Flat and elongated lipid-like densities (green) close to W939 in cryo-EM maps of detergent-solubilized homologues of hATP7B.**

A) Density next to W939 of DDM/CHS-solubilized and GDN-purified inactive hATP7B from HEK-F.

B) Density next to W939 of LMNG/CHS-solubilized and GDN-purified *Xt*ATP7B from *Pichia pastoris*.

C) Density next to W561 of DDM/CHS-solubilized and LMNG/GDN/CHS-purified *Os*HMA4 from *Saccharomyces cerevisiae*.

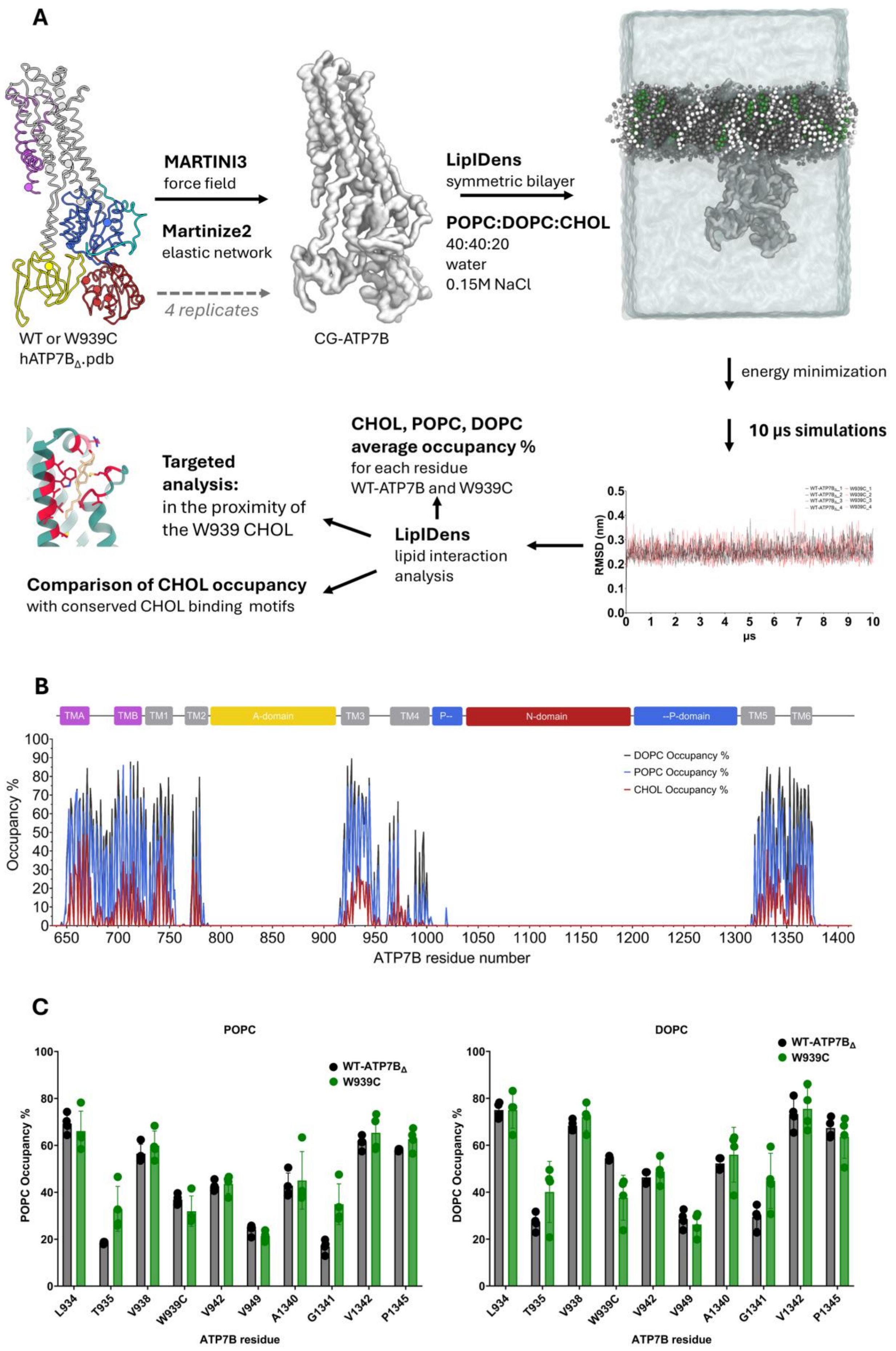


**Suppl. Fig. 12: MD Simulation workflow and condensed results.**

A) MD simulations and subsequent lipid occupancy analysis workflow. The WT-ATP7B$_{\Delta}$ or W939C mutant .pdb files were used as input for all replicates of the WT and W939C simulations. For each replicate, the structure was independently coarse-grained, embedded in a lipid bilayer simulation box. Following equilibration and energy minimization, four 10 μs simulations were performed for both WT and W939C. Per-residue lipid occupancies were analysed for each simulation.

B) Per-residue CHOL, DOPC, POPC occupancy (percentage of simulation frames in which a certain lipid contacts a residue) for WT-ATP7B$_{\Delta}$. The two-dimensional topology of hATP7B$_{\Delta}$ is shown above the graph, aligned with the sequence of ATP7B.

C) Comparison of mean POPC, DOPC occupancy between WT and W939C ATP7B$_{\Delta}$ for residues surrounding the lipid density observed near W939 in the cryo-EM map. Data from four independent 10 μs simulations with the WT-ATP7B$_{\Delta}$ and the W939C-ATP7B inputs. Data are presented as mean±SD. No significant differences between WT and W939C occupancies were observed for POPC and DOPC after multiple Welch's t-tests followed by Benjamini-Hochberg false discovery rate correction at 5%.

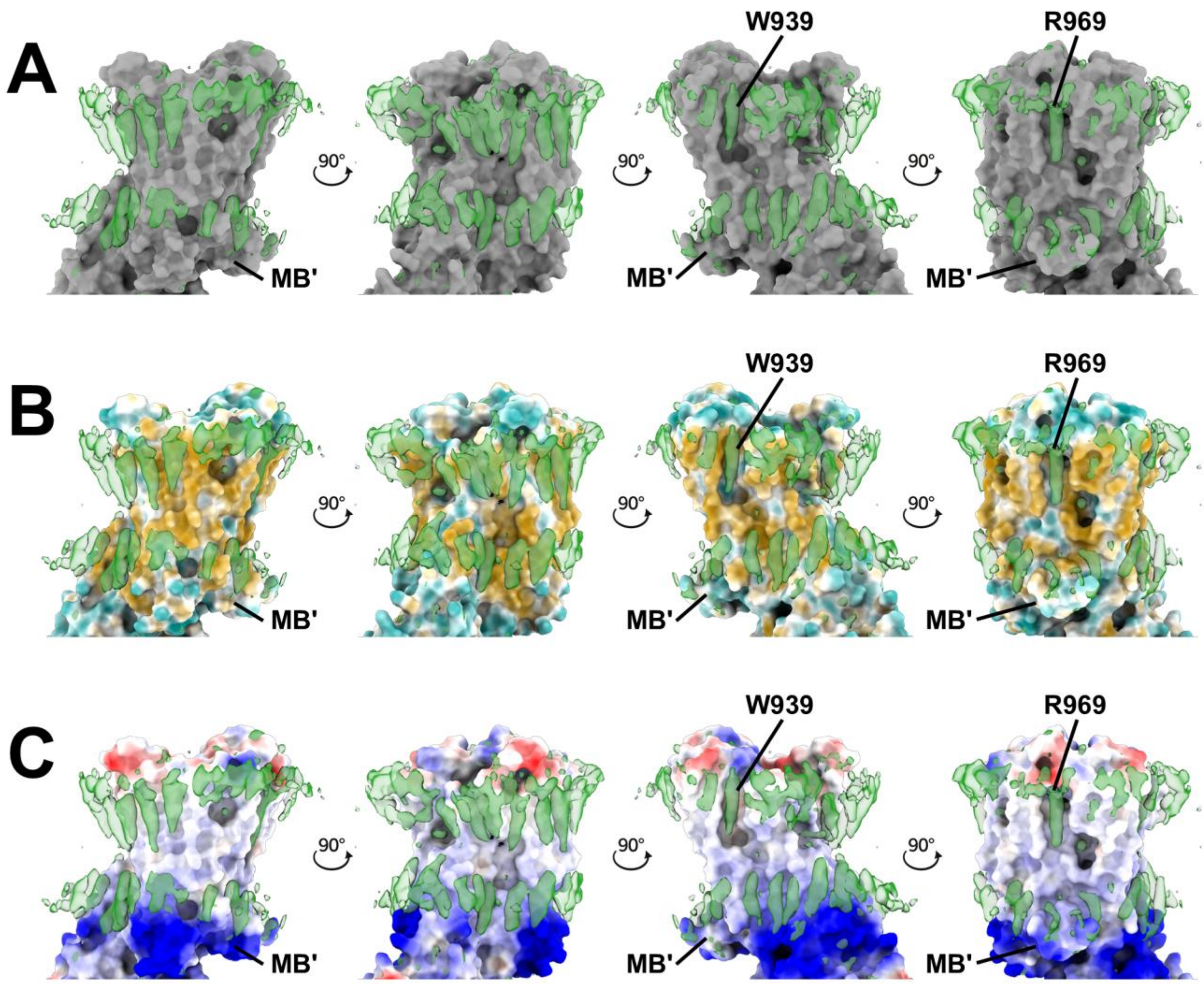


**Suppl. Fig. 13: MSP1E3D1-ATP7B$_{\Delta}$ surface representation with lipid densities from the EM map visualized at 9.2 sigma. Lipid-like densities adjacent to the WD mutation sites W939 and R969, which feature distinct pockets, are highlighted, with the MB' helix labelled for reference.**

A) Solid grey surface.
B) Hydrophobic surface. The molecular surface was colored according to the Fauchère molecular lipophilicity potential (MLP), with dark goldenrod indicating lipophilic and dark cyan indicating hydrophilic regions.
C) Electrostatic surface. The molecular surface was coloured according to Coulombic electrostatic potential using a red-white-blue colour scale (-10 to +10 $kcal \cdot mol^{-1} \cdot e^{-1}$), with blue representing positive electrostatic potential and red representing negative electrostatic potential.

**Supplementary Video 1**

WT-ATP7B MD simulation snapshot – CHOL near the W939 residue

**Supplementary Table 1:** Cryo-EM data collection, refinement, and validation statistics.

| Model | $ATP7B_{\Delta}$ | MBD5-up | MBD5-down |
|---|---|---|---|
| PDB | 50PI | 50PJ | 50PK |
| EMDB | 100822 | 100823 | 100824 |
| ***Data collection*** | | | |
| *Microscope* | Titan Krios | Titan Krios | Titan Krios |
| *Detector* | Gatan K3 with BioQuantum energy filter | Gatan K3 with BioQuantum energy filter | Gatan K3 with BioQuantum energy filter |
| *$C_s$ (mm)* | 2.7 | 2.7 | 2.7 |
| *Magnification* | 130,000x | 130,000x | 130,000x |
| *Voltage (kV)* | 300 | 300 | 300 |
| *Electron exposure ($e^-/Å^2$)* | 50 | 50 | 50 |
| *Defocus range (μm)* | 0.8 to 2.0 | 0.8 to 2.0 | 0.8 to 2.0 |
| *Pixel size (Å)* | 0.645 | 0.645 | 0.645 |
| *Micrographs collected* | 3,876 | 3,876 | 3,876 |
| *Micrographs used* | 2,913 | 2,913 | 2,913 |
| ***Reconstruction*** | | | |
| *Software* | cryoSPARC v4.6-4.7.0 | cryoSPARC v4.6-4.7.0 | cryoSPARC v4.6-4.7.0 |
| *Initial particle images (no.)* | 988,320 | 988,320 | 988,320 |
| *Final particle images (no.)* | 150,030 | 15,608 | 12,960 |
| *Symmetry imposed* | C1 | C1 | C1 |
| *Map global resolution (Å)* | 2.84 | 4.49 | 4.16 |
| *FSC threshold* | 0.143 | 0.143 | 0.143 |
| *Map local resolution range ($25^{th}$ to $75^{th}$ percentile; Å)* | 2.49 to 3.70 | 6.32 to 9.09 | 6.77 to 10.52 |
| ***Refinement*** | | | |
| *Software* | Isolde & Phenix real-space refinement | Isolde & Phenix real-space refinement | Isolde & Phenix real-space refinement |
| *Initial model used* | AlphaFold 3 prediction | AlphaFold 3 predictions & $ATP7B_{\Delta}$ | AlphaFold 3 predictions & $ATP7B_{\Delta}$ |
| *Map Sharpening B factor ($Å^2$)* | N/A (unsharpened map) | N/A (unsharpened map) | N/A (unsharpened map) |
| *Model resolution, d_model (Å)* | 3.2 | 4.6 | 4.2 |
| *Model-to-map FSC resolution (Å), FSC = 0.5* | 3.0 | 7.4 | 7.1 |
| *Model-to-map CC (mask)* | 0.83 | 0.71 | 0.71 |
| ***Model Composition*** | | | |
| *Non-hydrogen atoms (no.)* | 5,889 | 6,747 | 6,796 |
| *Protein residues (no.)* | 739 | 893 | 898 |
| *Ligands* | 1 MG, 1 ALF, 11 CLR | 1 MG, 1 ALF | 1 MG, 1 ALF |
| ***B factors ($Å^2$)*** | | | |
| *Protein* | 126.03 | 276.64 | 220.35 |
| *Ligands* | 124.98 | 45.80 | 45.80 |
| ***R.m.s. deviations*** | | | |
| *Bond lengths (Å)* | 0.006 | 0.014 | 0.006 |
| *Bond angles (°)* | 1.503 | 1.812 | 1.466 |
| ***Validation*** | | | |
| *MolProbity score* | 1.12 | 0.98 | 1.05 |
| *Clashscore* | 3.27 | 2.11 | 2.60 |
| *Rotamers outliers (%)* | 0.65 | 0.54 | 0.94 |
| *CaBLAM outliers (%)* | 0.27 | 1.02 | 0.34 |
| *Cβ outliers (%)* | 0.00 | 0.00 | 0.00 |
| ***Ramachandran plot*** | | | |
| *Favoured (%)* | 98.78 | 98.54 | 98.88 |
| *Allowed (%)* | 1.22 | 1.46 | 1.12 |
| *Outliers (%)* | 0.00 | 0.00 | 0.00 |

**Supplementary Table 2:** Identifiers for Wilson Disease-associated missense variants mentioned in the manuscript.

| ATP7B variant | mRNA transcript NM_000053.4 | dbSNP ID | ClinVar ID |
|---|---|---|---|
| R778L | c.2333G>T | *rs28942074* | 3852 |
| R778W | c.2332C>T | *rs137853284* | 456553 |
| R778G | c.2332C>G | *rs137853284* | 156283 |
| T935M | c.2804C>T | *rs750019452* | 188713 |
| W939C | c.2817G>C | *rs1057517310* | 1409871 |
| W939C | c.2817G>T | *rs1057517310* | 371483 |
| G943D | c.2828G>A | *rs779323689* | 189058 |
| G943R | c.2827G>C | *rs28942076* | 3072174 |
| G943S | c.2827G>A | *rs28942076* | 3856 |
| R969Q | c.2906G>A | *rs121907996* | 3860 |
| H1069Q | c.3207C>A | *rs76151636* | 3848 |
| G1341D | c.4022G>A | *rs779494870* | 665925 |
| G1341V | c.4022G>T | *rs779494870* | - |
| G1405S | c.4213G>A | *rs189601972* | 444316 |

**Supplementary Table 3: Truncations and WD-associated variants of the ATP7B C-terminal tail, before and after residues 1404-1410 at the A-, N-domain and MBD5 junction.** *In vitro* studies showed that variants and deletions affecting the proximal, structured portion of the C-terminal tail, extending approximately to W1410, tend to compromise protein activity and stability, whereas more distal variants more frequently affect copper-responsive trafficking [PMID:14962673, 37660282, 24831241, 35245129], although this distinction is not absolute and may depend on cellular context [PMID:32778786].

| ATP7B variant | mRNA transcript NM_000053.4 | dbSNP ID | ClinVar ID | Clinical evidence |
|---|---|---|---|---|
| E1382Ter | c.4144G>T | *rs213841473* | 1162217 | Pathogenic/Likely pathogenic |
| Y1384fs | c.4145_4148dup | *rs254754383* | 2763268 | Pathogenic |
| E1385Ter | c.4152dup | *rs2547543720* | 2020602 | Pathogenic/Likely pathogenic |
| A1388fs | c.4162del | — | — | Reported in WD patients (PMID: 34470610) |
| M1392K | c.4175T>A | — | — | Reported in WD patients (PMID: 34470610,20931554) |
| Q1399Ter | c.4195C>T | *rs2138407151* | 1456240 | Pathogenic/Likely pathogenic |
| Q1399fs (*) | c.4195del | *rs886041336* | 280040 | Pathogenic/Likely pathogenic |
| V1400fs | c.4198del | *rs2547542768* | — | Pathogenic/Likely pathogenic |
| G1405S | c.4213G>A | *rs189601972* | 444316 | Likely pathogenic/Uncertain significance |
| W1410Ter | c.4229G>A | *rs2547542223* | 1929153 | Pathogenic/Likely pathogenic |
| W1410Ter | c.4230G>A | *rs2547542207* | — | Pathogenic |
| S1413fs | c.4238_4245del | *rs2547541937* | — | Pathogenic |
| R1415fs | c.4242del | *rs1057517191* | — | Likely pathogenic |
| D1420fs | c.4258dup | *rs1956866547* | — | Pathogenic |
| Y1424Ter | c.4272T>G | — | — | Reported in WD patients (PMID: 34470610) |
| S1429fs | c.4283dup | *rs155528219* | 551073 | Pathogenic/Likely pathogenic |
| T1434fs | c.4299del | *rs254754100* | 2166696 | Pathogenic |

| | | | | |
|---|---|---|---|---|
| A1445P | c.4333G>C | — | — | Reported in WD patients (PMID: 34470610,20931554) |
| W1452Ter | c.4355G>A | *rs780486131* | 2846900 | Pathogenic/Likely pathogenic |
| W1452Ter | c.4356G>A | *rs134117218* | | Pathogenic/Likely pathogenic |
| S1453fs | c.4354_4357dup | *rs254753969* | 2161078 | Pathogenic |
| G1458fs | c.4373del | *rs2547539500* | | Pathogenic |
| R1459fs | c.4374_4375del | *rs2138386032* | 1071568 | Pathogenic |

*(*) The ATP7B c.4195del variant (p.Gln1399ArgfsTer6) corresponds to the Saudi founder frameshift variant historically reported as 4193delC/Q1399R [PMID: 11054498, 14748773]. During our literature review, we noted that c.4196A>G (p.Gln1399Arg) has been reported as a Wilson disease-associated ATP7B variant in several secondary sources [PMID: 25496901, 39767741, 38397079, 35208595, 35245129]. However, the primary Saudi studies cited in support of this association describe the distinct frameshift variant c.4195del (historically 4193delC), rather than the c.4196A>G missense variant. We were unable to identify an independent primary report supporting an association of c.4196A>G (p.Gln1399Arg) with Wilson disease.*

**Supplementary Table 4:** Primers used in this study. Mutated bases are underlined.

| Primer name | Primer sequence (5'→3'), mutation underlined |
|---|---|
| E754K mut fwd | GTTGCGGTTGCTAAAAAGGCTGAAAGATCC |
| E754K mut rev | GGATCTTTCAGCCTTTTTAGCAACCGCAAC |
| W939C mut fwd | CCTTGACTTTGGTCGTCTGTATCGTCATTGGTTTCATCG |
| W939C mut rev | CGATGAAACCAATGACGATACAGACGACCAAAGTCAAGG |
| R969E mut fwd | CTGAAGTCATTATCGAATTTGCTTTCCAAAC |
| R969E mut rev | GTTTGGAAAGCAAATTCGATAATGACTTCAG |
| R969Q mut fwd | CTCCCAAACTGAAGTCATTATCCAATTTGCTTTCCAAACTTCC |
| R969Q mut rev | GGAAGTTTGGAAAGCAAATTGGATAATGACTTCAGTTTGGGAG |
| D1027N mut fwd | GATCAAGACTGTTATGTTCAACAAGACCGGTACTATCAC |
| D1027N mut rev | GTGATAGTACCGGTCTTGTTGAACATAACAGTCTTGATC |
| H1069Q mut fwd | GAAGCCTCCTCTGAACAACCATTGGGTGTTGC |
| H1069Q mut rev | GCAACACCCAATGGTTGTTCAGAGGAGGCTTC |
